# “Photosynthetic and Genetic Adaptations Underpinning the Resilience of *Cistanthe longiscapa* in the Atacama Desert”

**DOI:** 10.1101/2024.12.08.627406

**Authors:** Omar Sandoval-Ibáñez, Patricio Tapia-Reyes, Aníbal Riveros, Ricardo Yusta, Shengxin Chang, Paulina Ossa, Ricardo Nilo-Poyanco, Adrián A. Moreno, Alvaro Miquel, Andrea Miyasaka Almeida, Andrés Zurita-Silva, Daniela Orellana, Carlos Baeza, Francisca Blanco-Herrera, Alex Di Genova, Miguel L. Allende, Mauricio González, Alejandro Maass, Martin Montecino, Rodrigo A. Gutiérrez, Ralph Bock, Claudio Meneses, Ariel Orellana

## Abstract

- The Atacama Desert is one of the most hostile environments for life. However, the plant species *Cistanthe longiscapa* (C. longiscapa) completes its life cycle in the Atacama Desert after sporadic rainfall.
- Physiological analyses under controlled environmental conditions revealed superior photosynthetic performance, better light acclimation mechanisms, and larger accumulation of photosystem II in *C. longiscapa* compared to its mesophilic sister species.
- *C. longiscapa* shows evolutionary expansions in gene families related to DNA repair, photosynthesis, and protein homeostasis. In addition, we observed substantial gene duplication and polymorphic variations between coastal and inland populations in the Atacama Desert. Finally, our assembled mitochondrial genome provides genetic information for all DNA-containing compartments of *C. longiscapa*.
- Diurnal oscillations of malic acid and time-resolved transcriptome analyses of plants harvested in the Atacama Desert indicate that *C. longiscapa* engages in CAM metabolism. We observed significant differences in transcripts encoding plastid-localized proteins, including those involved in carbon metabolism, light harvesting, and photoprotection, highlighting the critical role of chloroplasts in the adaptation of *C. longiscapa* to the Atacama Desert.
- Our study provides physiological and genetic evidence for the adaptations of *C. longiscapa* and advances our understanding of how plants can cope with extreme environmental conditions.

## Introduction

Climate change is a major threat to sustainable crop production an is driving research towards a better understanding of plant responses to extreme suboptimal conditions (Eckardt *et al*., 2022). As sessile organisms, land plants must cope with stresses throughout their life cycle, including drought, heat and high light stress (Hu *et al*., 2020; Song *et al*., 2021). Plastids are particularly sensitive to abiotic changes, and must rapidly adapt the photosynthetic capacity to suboptimal conditions. Therefore, photosynthesis has been the main target for plant breeding, aimed at increasing plant resilience to suboptimal environmental conditions while maintaining biomass production (Croce *et al*., 2024). In land plants, photosynthesis comprises two reactions: the Calvin-Benson-Bassham (CBB) cycle, which is responsible for the CO_2_ fixation, and the electron transport chain (ETC), mediated by the thylakoid-anchored photosynthetic complexes, that supply ATP and NADPH to the CBB cycle (Johnson, 2016). While three mechanisms for CO_2_ assimilation are described, known as C_3_, C_4_, and crassulacean acid metabolism (CAM), CAM photosynthesis has been studied for decades, because it increases the water use under extreme drought conditions. CAM performing species acquire CO₂ at night, which is incorporated to phosphoenolpyruvate (PEP) via PEP carboxylase (PEPC) and finally to malate by two consecutive reactions (Winter & Smith, 2022). During the day, stomata remain closed, and malate acts as an organic acid carrier that delivers CO₂ to Rubisco (Winter & Smith, 2022). While the origin of CAM photosynthesis remains unclear, recent studies suggest a continuous evolutionary model from C_3_ to CAM (Bräutigam *et al*., 2017), with species using C_3_ - CAM pathways for carbon capture (known as CAM facultative), and plants relying exclusively on CAM as the primary carbon fixation mechanism. In particular, CAM photosynthesis has been studied for decades, due to its ability to increase water use under extreme drought conditions (Heyduk, 2022). However, engineering C_3_ plants to use CAM remains challenging due to the lack of sufficient genetic and physiological information on plants that perform constitutive CAM, making it difficult to distinguish metabolic from stress-related responses (Tan & Chen, 2023). In addition to carbon fixation pathways, plants evolve various photoprotective mechanisms to avoid the production of reactive oxygen species (ROS) during the ETC under stress conditions. For example, the production of photoprotective pigments (Simkin *et al*., 2022), the protection of photosynthetic complexes involved in ETC (Lei Li *et al*., 2018) and mechanisms of light maintain photosynthesis. However, it is still not clear how the photosynthesis of extremophile plants has adapted (Fernández-Marín *et al*., 2020).

The combination of intense UV radiation and minimal precipitation makes the Atacama Desert, located in northern Chile, one of the most arid and inhospitable regions on Earth (Cordero *et al*., 2018) and an ideal setting to study mechanisms of plant resilience. Among the sparse flora adapted to this harsh environment, *Cistanthe longiscapa* (*C. longiscapa*) stands out as a particularly resilient species (Holtum *et al*., 2021). The seeds of *C. longiscapa* remain dormant for years and germinate after sporadic rainfall, completing a full life cycle in 2-3 months even long after the available water has evaporated. *C. longiscapa* belongs to the Montiaceae family within the Caryophyllales order, and is the main plant species contributing to the phenomenon known as the “flowering desert”. *C. longiscapa* performs CAM photosynthesis even under unrestricted water conditions (Holtum *et al*., 2021), making it an excellent plant model for studying the mechanisms of photosynthesis.

In this work we investigate the physiological and genetic advantages of *C. longiscapa* in order to understand the mechanisms of plant adaptation that facilitate survival in the Atacama Desert. We compared the photosynthetic performance of *C. longiscapa* with that of the related Caryophyllales species *Amaranthus hypochondriacus* and *Cistanthe grandiflora* (Lightfoot *et al*., 2017; Yao *et al*., 2019; Netshimbupfe *et al*., 2022) to gain insights into the adaptation mechanisms that enable *C. longiscapa* to cope with extreme light intensities. In addition, we report the sequence of the *C. longiscapa* nuclear and mitochondrial genomes and explore the existing nuclear genome variation across populations. Finally, we provide a comprehensive transcriptomic analysis and associated gene expression profiles of *C. longiscapa* under Atacama Desert conditions. By integrating physiological and -omics data, our study provides a comprehensive understanding of the adaptive strategies employed by a desert plant and a rich resource for future research on plant resilience in extreme habitats.

## Materials and Methods

### Plant material, sample collection and *in vitro* growth conditions

The plant species used in this study were *Cistanthe longiscapa*, *Amaranthus hypochondriacus* cultivar Router Dom, *Cistanthe grandiflora*, and *Arabidopsis thaliana* ecotype Columbia-0. Samples and seeds of *Cistanthe longiscapa* were collected from five different Locations (Table S1). Seeds of *Cistanthe grandiflora* were obtained from the Instituto de Investigaciones Agropecuarias (INIA) under stock NAT1430-LP19. Seeds of *Amaranthus hypochondriacus* were obtained from the Leibniz Institute of Plant Genetics and Crop Plant Research (IPK) Gatersleben under accession number AMA 20. *C. longiscapa* plants were collected from different locations (Table S1). Plants from location 1 were used for genome assembly. Plants collected from locations 2, 3, and 5 were used for genome diversity analyses and mitochondrial genome assembly. *C. longiscapa* plants collected from location 4 were used for transcript analyses of leaves harvested every 6 hours over a 24 time-course (first harvest at 7 am and last harvest at 1 am of the next day) and tissue-specific libraries for transcript enrichment (roots, succulent stem, leaf, inflorescence stem and flowers).

For seed propagation and *in vitro* cultivation, *C. longiscapa*, *A. hypochondriacus* and *C. grandiflora* seeds were surface sterilized with a solution of 1% sodium hypochlorite solution, washed with sterile water, and stratified for 3-5 days. For sterile culture, seeds were germinated in 0.5x Murashige & Skoog medium supplemented with 1% sucrose. Plants were grown at 22°C, under a light intensity of 100 μmol photons m^−2^s^−1^, and a light-dark cycle of 16 h/8 h.

### Imaging Pulse-Amplitude Modulation (Imaging-PAM) analysis

*C. longiscapa*, *A. hypochondriacus* and *C. grandiflora* plants grown under sterile conditions for 3 weeks were dark-adapted for 30 min prior to PAM measurements, using the IMAGING-PAM M-Series. Induction curves and recovery for *C. longiscapa*, *A. hypochondriacus* and *C. grandiflora* were performed at light intensities ranging from 33 to 1,160 μmol photons m^−2^ s^−1^. Data collection setting are listed in Methods S1.

### Thylakoid isolation, PAA gel electrophoresis and immunodetection

Thylakoid isolation was performed according to Pribil et al. (Pribil *et al*., 2018). Briefly, 3-week-old leaves of *C. longiscapa*, *A. hypochondriacus* and *C. grandiflora* plants were ground in liquid nitrogen, resuspended in grinding buffer [50 mM HEPES KOH pH7.5; 330 mM sorbitol; 2 mM EDTA; 1 mM MgCl_2_; 5 mM ascorbate; 0.05% bovine serum albumin] and filtered through two layers of Miracloth ®. The samples were centrifuged at 6,000g for 4 minutes at 4°C and the pellet was resuspended in shock buffer [50 mM HEPES KOH pH7.5; 5 mM sorbitol; 5 mM MgCl_2_]. The samples were then centrifuged at 6,000g for 4 min at 4°C, washed once with storage buffer [50 mM HEPES KOH pH7.5; 100 mM sorbitol; 10 mM MgCl_2_], and finally resuspended in a small volume of storage buffer. Chlorophyll content was quantified according to Porra et al. 1989 (Porra *et al*., 1989).

Total protein samples or isolated thylakoids were solubilized in Laemmli buffer and resolved in a 12.5 % or 15 % SDS-PAGE gel. Immunodetection was performed using the antibodies listed in Table S2. BN-PAGE and 2-dimensional SDS-PAGE were performed according to Jarvi et al. (Jarvi *et al*., 2011), with the following modifications. Thylakoid samples were solubilized in 1% β-DDM for 2 min at 4°C and resolved in an 8-13.5 % PAA gradient (32:1 acrylamide:bisacrylamide). For second-dimension SDS-PAGE, the firt dimension gel strips were solubilized with Laemmli buffer [138 mM Tris HCl, pH6.8; 6 M urea, 22.2% glycerol, 4.3% SDS and 100 mM DTT] and loaded onto a 12.5% PAA gel. Immunoblots were performed on PVDF membranes (Sandoval-Ibáñez *et al*., 2022).

### Chlorophyll *a* fluorescence emission at low temperature (77 °K)

The kinetic movement of the antenna (state transitions) was analyzed according to Pribil et al. (Pribil *et al*., 2018), with the following modifications. *C. longiscapa* and *C. grandiflora* plants were illuminated with far-red light (730 nm) for 2 h, and then exposed to darkness for 5 and 10 min. At each time point, samples were collected, ground in liquid nitrogen, resuspended in a buffer containing 50 mM HEPES pH 7.5, 330 mM sorbitol and 1 mM MgCl_2_, and the chlorophyll concentration was adjusted to a minimum and a maximum of 8 μg and 10 μg of chlorophyll mL^−1^, respectively. Low temperature chlorophyll *a* fluorescence emission spectra at 77 °K were recorded using an F-6500 fluorometer (Jasco Inc.), with an excitation wavelength of 430 nm and a bandwidth of 10 nm in the range of 650-780 nm in 0.2 nm intervals. The emission spectra were normalized to the emission values at 688.5 nm.

### D1 degradation kinetics

The D1 degradation kinetic were performed as described by Pribil et al. (Pribil *et al*., 2018). Briefly, 3-week-old *C. longiscapa* and *C. grandiflora* plants were vacuum infiltrated with a 10 mM lincomycin solution three times for 10 min, and incubated overnight at 4°C in the dark. The plants were then exposed to light at an intensity of 800 μmol photons m^−2^ s^−1^ for 0, 0.5, 1, 2, 3 and 4 h. At each time point, the plants were dark adapted for 10 min and Fv/Fm was recorded as an indicator of PSII activity.

### Genome size determination

For flow cytometry-based genome size estimation, 100 mg of *A. thaliana* or *C. longiscapa* leaves were minced at 4°C in 2 mL of a buffer containing 45 mM MgCl_2_, 30 mM sodium citrate, 0.1% Triton X-100 and 20 mM MOPS pH 7.0 (Galbraith *et al*., 1983; Zonneveld, 2021). Cell debris was removed by filtration through a 70 μm filter, and the filtrate was incubated with 50 µg mL^−1^ of RNase A and 50 µg mL^−1^ of propidium iodide (PI) at 4°C for 30 min. Flow cytometry was performed on a BD FACSAria ™ III with a flow rate of 200 µL min^−1^, laser excitation at 488 nm, and PI detection filters 585/42 (FL-2) and 616/23 (FL-3). *A thaliana* was used as a reference, for genome size estimation, using the calculation method described by Al-Qurainy et al. (Al-Qurainy *et al*., 2021).

### Karyotype determination

For chromosome preparation, *C. longiscapa* was grown on water–moistened filter paper at 24°C. Roots from seedlings were excised and pre–treated with 8–hydroxyquinoline solution (2 mM aqueous solution) at 4°C for 24 h, fixed in a freshly prepared mixture of absolute ethanol/glacial acetic acid (3:1) for 24 h and stored in 70% ethanol at –20°C. To determine the chromosome number, the roots were stained with 0.1% orcein, and the chromosomes were prepared by squashing the roots tips (Baeza *et al*., 2007).

### Nucleic acid extraction

For genome assembly and Illumina NovaSeq6000 sequencing, high molecular weight genomic DNA was extracted by Polar Genomics (http://polargenomics.com) (Zhang *et al*., 1995) from a single plant from location 1 (Table S1).

For genetic diversity analysis and mitochondrial genome assembly, high-quality DNA was prepared from frozen leaf material of *C. longiscapa* harvested from locations 2, 3 and 5. Cuticles were removed with a scalpel prior to grinding in liquid nitrogen, and the ground tissue was lyophilized until the sample was completely dry for 24 hours. A total of 1 mg of lyophilized tissue was processed with the MasterPure Complete DNA and RNA purification kit (Illumina) according to the protocol provided by the supplier. DNA integrity was verified by gel electrophoresis, and total DNA was quantified by fluorometry using the Qubit dsDNA HS assay (Invitrogen).

For total RNA extraction, plant samples were ground in liquid nitrogen and the powder was resuspended in NTES buffer [20mM Tris pH 8.0, 100mN NaCl, 10mM EDTA pH 8.0 and 1% SDS]. The samples were mixed with 2 volumes of phenol:chloroform (1:1). After centrifugation, the aqueous phase was separated, and the total RNA was extracted from the aqueous phase was extracted with TriZol ®. RNA quality was determined by capillary electrophoresis (Fragment Analyzer, Agilent) and by fluorometry using Qubit (Invitrogen).

### DNA and RNA sequencing

Genomic DNA was prepared in SMRTbell templates (Pacific Biosciences, part no. 100-259-100), SMRTbells were bound to the v2.0 polymerase and sequenced on the PacBio RS II system using 10 hour movies, and data were collected from a total of 25 SMRT Cell 1M. For Illumina NovaSeq6000 sequencing, genomic DNA was sonicated, and the library was generated using the TruSeq DNA Nano Kit (Illumina). The library was sequenced using the HiSeq4000, 150 bp PE (Table S3).

In order to facilitate genome annotation, RNA samples were obtained from five distinct plant tissues at location 4 and six different tissues from the same individual plant used for genome annotation collected at location 1 were used (Table S1). The libraries were generated by the TruSeq RNA Sample Preparation v2® (Illumina) (Table S3), and sequenced using the Illumina Hiseq 2500 platform (Macrogen, Seul, Korea).

For the construction of the ISO-Seq libraries, equal content of RNA extracted from the six tissues of the individual plant from Location 1 were pooled into a single sample. The pooled RNA was retrotranscribed, and prepared into SMRTbell templates (Pacific Biosciences, part no. 100-259-100) (Table S3). The libraries were sequenced on the PacBio RS II system using 10-hour movies. Data from two SMRT Cell 1M were collected. For transcriptomic analysis, three independent pools of leaf samples of plants were collected from Location 4 during a six-hour time course. Twelve libraries were generated by the TruSeq RNA Sample Preparation v2® (Illumina) (Table S3), and were sequenced using the HiSeq4000 platform (Macrogen, Seoul, Korea).

### Genome assembly of *C. longiscapa*

To assemble a pre-processed library based on Illumina reads, the Genome_NS_T01 library was trimmed by removing low quality 3’ borders, sequencing adaptors and reads with less than 35 bp (Table S4). Plastid reads were removed from the trimmed library by alignment to the *C. longiscapa* reference chloroplast genome (Stoll *et al*., 2017a). The unaligned reads were saved as a fastq file for the hybrid correction of the PacBio libraries and assembly of a hybrid reference genome.

To assemble a hybrid reference genome, the 25 raw PacBio libraries obtained from *C. longiscapa* were concatenated in a single Fasta file. The resulting file was assembled and combined with the plastid-free Illumina pre-processed reads. The resulting scaffold reference genome was post-refined in two steps. First, the gaps were removed using the corrected PacBio data as input. Then, the erroneous consensus base pairs were corrected to produce a refined version of the reference genome scaffold. The refined genome scaffold was rearranged using the genome assembly of *Portulaca amilis* (Gilman *et al*., 2022), *Selenicereus undatus* (Zheng *et al*., 2021) and *Talinum fruticosum* (NCBI Accession: PRJNA659383; ID: 659383) as references, obtaining the final version of the reference genome. To validate the genome assembly and scaffolding, BUSCO v.5.2.2 software (Manni *et al*., 2021) was used to analyze the completeness of the PacBio raw data, the long-read primary assembly, and the hybrid scaffolding. Finally, each library was aligned to the genome scaffold. See Methods S2 for further information.

### Genome annotation of *C. longiscapa*

The annotation of the *C. longiscapa* genome was performed with the version of SwissProt-uniprot viridiplantae retrieved in December 2021 as a protein database, in addition to the complete proteome of 15 *Caryophyllales* genomes (Table S5). The completeness of the annotation with the software BUSCO v.5.2.2 (Manni *et al*., 2021). Functional annotation was performed by BLASTp alignment of predicted proteins-coding genes to the SwissProt-viriplantae database, and InterProScan to Pfam database. To complement the annotation, we also used the online platforms Mercator4/Mapman (https://plabipd.de/portal/mercator4, November 2022), eGGNog mapper v.2 with eGGNog v.5.0 Database (http://eggnog5.embl.de/#/app/home, July 2022) and KASS (https://www.genome.jp/kegg/kaas/, November 2022). See Methods S3 for further information.

### Comparative genomics

To build the ultrametric tree, the proteomes of 15 Caryophyllales species were obtained from different repositories (Table S5). The sequences from the repositories, and the sequences from *A. thaliana* and *C. longiscapa* were used as input, using the calibration points for *A. thaliana* – *Fagopyrum tataricum* (125 MYA), *Portulaca amilis* – *Chenopodium quinoa* (90 MYA), and *Fagopyrum tataricum* – *Beta vulgaris* (106 MYA). The expansion/contraction of orthologous groups was calculated using the software Cafe v.5.0.0 (Mendes *et al*., 2021). For more information, see Methods S4.

### Short variant calling

For short variant discovery, the 12 WGS libraries from three different locations (location 2, 3 and 5) were normalized to the library with the lower number of reads. The uBAM files from the normalized libraries were used for short variant discovery (SVD) which was performed with GATK v.4.0.9.0 (van der Auwera *et al*., 2013) using the lastest scaffolded version of the reference genome. The final 12 VCF files (one for each library) were filtered for QUAL > 100. To obtain the unique or shared SNPs for each VCF file, we used the “isec” package from bcftools v.1.9 (Danecek *et al*., 2021). See Methods S5 for more information.

### Draft assembly of the mitochondrial genome

Raw data from five libraries of short Illumina reads from location 5 were used to assemble the mitochondrial genome. Annotation of protein-coding genes and rRNAs was performed in the online version of GeSeq v2.03 (Tillich *et al*., 2017) using the Caryophyllales mitochondrial genes as a reference, and later were manually verified. For tRNA gene, we used the GeSeq built-in tRNAscan-SE v2.0.7 (Chan *et al*., 2021). Maps of the assembled mitochondrial DNA circles were drawn with the OGDRAW tool v1.1.1 (Lohse *et al*., 2007; Greiner *et al*., 2019). For more information, see Methods S6.

### Total acid and malic acid determination

For the determination of total acid, leaves were ground in liquid nitrogen, weighed and resuspended in water. Total acid was determined by titration to pH 7.0 using a 0.01N NaOH as previously described (KEELEY & KEELEY, 1989) . For the quantification of malic acid, leaves were ground in liquid nitrogen, weighed and resuspended in water. Insoluble material was removed by centrifugation and malic acid was separated by HPLC according to the procedure described by Tasev et al. (Tasev *et al*., 2016). The reverse-phase separation was performed on a C18 Aclaim 120 column (5 µm, 120 Å, 4.6 x 250 mm) under isocratic conditions at a flow rate of 1.0 mL/min. The mobile phase of 25 mM orthophosphoric acid (pH 2.5) was supplemented with 1% acetonitrile. The sample was injected into the HPLC system with a loop volume of 25 µL. Detection was carried out at a wavelength of 210 nm using a Thermo Ultimate 3000 diode array detector.

### Differential expression analysis and GO annotation

Reads were filtered and mapped to the reference genome. Reads with counts less than 5 were discarded, and the data were normalized to the libraries obtained at 7:00 using the median of ratios method. Sequences annotated as plastid- and mitochondrial-encoded transcripts were removed, and finally, the data were filtered by fold change (-1>FC>1), using an FDR <0.05. For GO enrichment annotation of consecutive time points, differentially expressed up- and down-regulated transcripts obtained in each pairwise comparison (T2/T1, T3/T2, T4/T3 and T1/T4) were analyzed with the Clusterprofiler v3.16.0 R package (Yu *et al*., 2012), using the Araport11 *Arabidopsis thaliana* annotation as input, and an adjusted *P*-value <0.05. See Methods S7 for further information.

## Results

### Description of Cistanthe longiscapa

*C. longiscapa* is the most iconic plant of the Atacama Desert and the main species during the desert bloom phenomenon (Fig.1). The leaves have succulent characteristics and make up the majority of the plant’s biomass (Fig S1a-d). The flowers have petals that account for the “blooming desert” phenomenon (Fig.1; Fig S1d,e). Successful laboratory propagation of *C. longiscapa* depends on cross-pollination (Fig. S1f-j). The seeds have a black coat and are approximately 1-2 mm in diameter (Fig. S1k). Stress conditions induce the accumulation of pigments exclusively in the leaf epidermis (Fig. S2a-b), corresponding to betalains (Fig. S2c).

**Fig.1:**
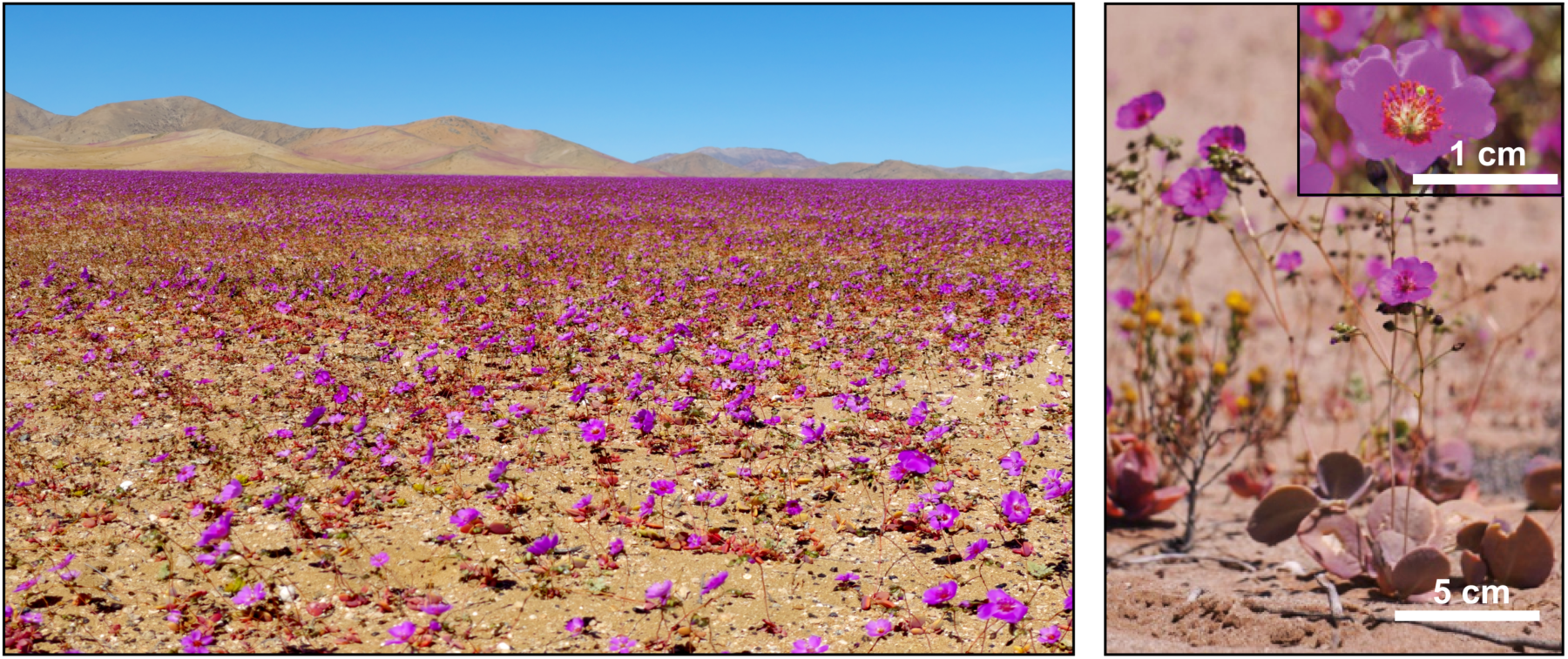
C. *longiscapa* is the most abundant plant in the Atacama Desert during the blooming episodes. Representative picture of *Cistanthe longiscapa* causing a desert bloom in the Atacama Desert. The right panel depicts a close-up of a group of plants (scale bar: 5 cm) and the inset shows the flower morphology (scale bar: 1 cm).

### *C. longiscapa* is adapted to high light intensities

The high solar radiation and low water availability of the Atacama Desert are two known factors that negatively affect photosynthesis (Croce *et al*., 2024). Nevertheless, *C. longiscapa* is able to complete its life cycle after occasional rainfall events, raising questions about the mechanisms of photosynthetic plasticity in *C. longiscapa*. We therefore investigated the photosynthetic performance of *C. longiscapa* using imaging-pulse amplitude modulation (PAM) fluorometry (Fig. 2). For comparison, a previously sequenced species in the order Caryophyllales, *Amaranthus hypochondriacus* (*A. hypochondriacus*) (Lightfoot *et al*., 2017) and *C. longiscapás* sister species, *Cistanthe grandiflora* (*C. grandiflora*), were included in these analyses. As the three species do not share a similar ecological niche, we used 3-week-old plants grown on sucrose-containing medium to minimize the effects of cultivation conditions on photosynthesis (e.g., water availability, temperature, and soil nutrient composition). A significantly higher effective quantum yield of photosystem II (Φ_II_, reflecting the activity of photosystem II) was observed in *C. longiscapa* compared to *A. hypochondriacus* and *C. grandiflora* (Fig. 2a, e). Compared to *A. hypochondriacus*, *C. longiscapa* also showed a faster recovery of Φ_II_ in the dark phase (Fig. 2a). While non-photochemical quenching (NPQ), which reflects the dissipation of excess light energy as heat, was significantly higher in *A. hypochondriacus* (at light intensities greater than 55 μmol photons m^−2^ s^−1^) and *C. grandiflora* (at light intensities greater than 230 μmol photons m^−2^ s^−1^), the relaxation of NPQ in the dark was similar in the three species (Fig. 2b, f). Next, we evaluated the qL parameter (reflecting the fraction of PSII reaction centres that are ‘open’, with the primary quinone-type acceptor Q_A_ being in the oxidized state) in the three species (Fig. 2c, g). While qL values were similar in *C. longiscapa* and *A. hypochondriacus* under increasing light intensities (Fig. 2c), they were lower in *C. grandiflora* (Fig. 2g). However, compared to *A. hypochondriacus,* a faster recovery was observed in *C. longiscapa* at early times during the dark phase (Fig. 2c), suggesting a faster ‘closing’ of PSII reaction centers (associated with faster oxidation of plastoquinol) in the donor side of PSII. Finally, the electron transport rate (ETR) was significantly higher in *C. longiscapa* compared to *A. hypochondriacus* and *C. grandiflora* at high light intensities (between 230-920 μmol photons m^−2^ s^−1^ and 530-1,160 μmol photons m^−2^ s^−1^, respectively; Fig.2d, h). Taken together, these results indicate that *C. longiscapa* has a better photosynthetic performance around PSII compared to *A. hypochondriacus* and *C. grandiflora*.

**Fig.2:**
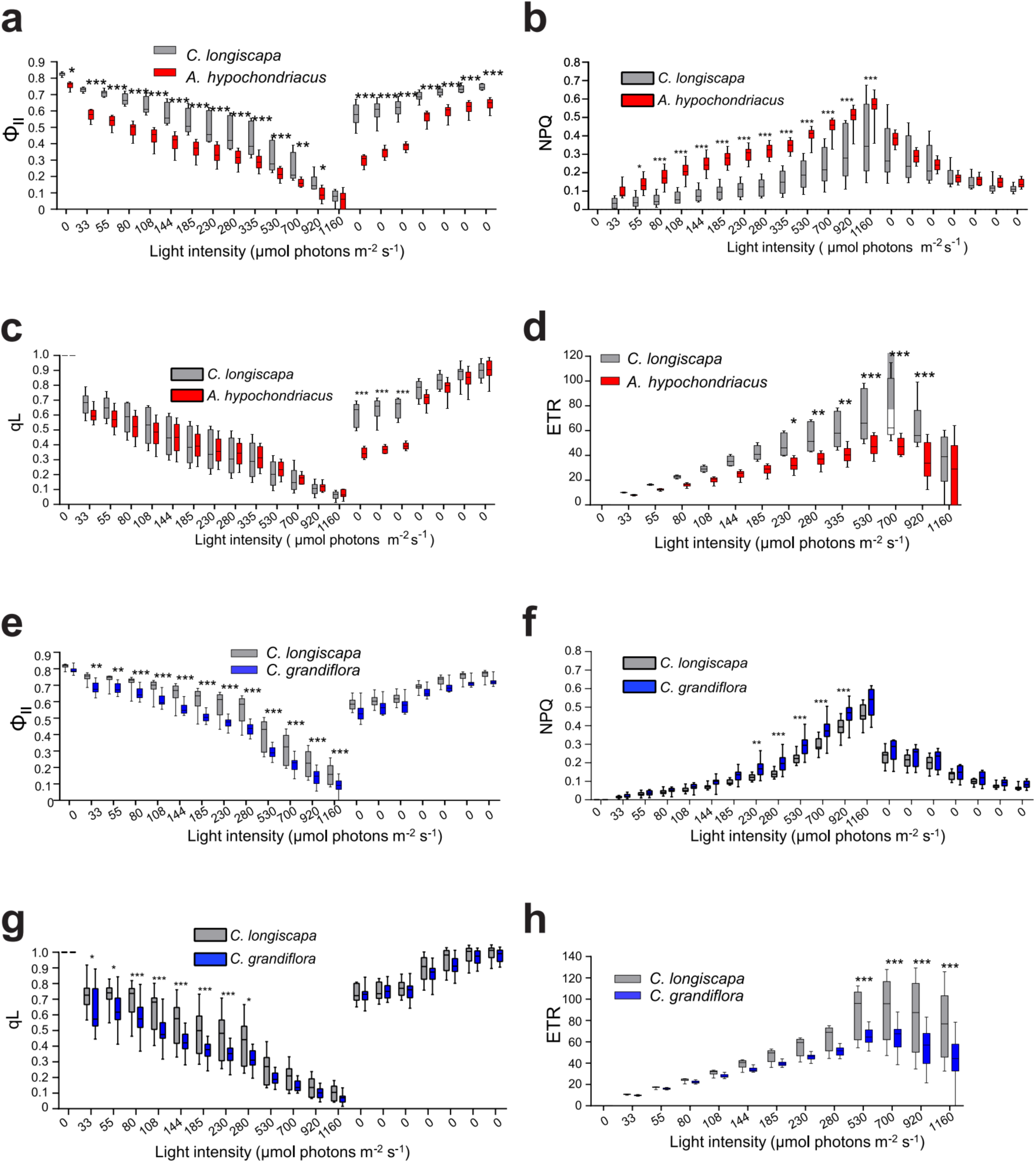
Photosynthetic performance of *C. longiscapa compared to A. hypochondriacus* and *C. grandiflora*. The chlorophyll *a* fluorescence parameters of (**a**, **e**) quantum yield of photosystem II (Φii), (**b, f**) non-photochemical quenching (NPQ), (**c**, **g**) qL and (**d, h**) electron transport rate (ETR) obtained by induction curves from *C. longiscapa* (grey) compared to *A. hypochondriacus* (red) and *C. grandiflora* (blue) grown under standard light conditions (100 μmol photons m^−2^ s^−1^) for 3 weeks are shown. Plants were dark adapted for 20 minutes prior to the measurement. The induction curve was recorded by increasing the light intensity from 0 to 1160 μmol photons m^−2^ s^−1^, followed by a recovery phase in the dark. n=10-15. Statistical significance was determined by two-way ANOVA, and *P*-values were adjusted with Sidak’s multiple comparison test comparing both species at the same light intensity. * *P*<0.05, ** *P*<0.01, *** *P*<0.001.

### *C. longiscapa* exhibited greater accumulation of PSII subunits and efficient light acclimation mechanisms

We then examined the accumulation of the main photosynthetic protein complexes in thylakoid samples isolated from *C. longiscapa* and *A. hypochondriacus* by SDS-polyacrylamide (PAA) gel electrophoresis and immunoblot analysis (Fig. 3a). While similar protein accumulation of PSI (as evidenced by analysis of the two diagnostic subunits PsaA and PSAD) and Cyt*b*_6_*f* (PetA, PetB) subunits were observed in *C. longiscapa* and *A. hypochondriacus*, a higher accumulation of diagnostic PSII subunits (PsbD and PsbA) was detected in *C. longiscapa* (Fig. 3a). Similarly, the accumulation of PSII core subunits was higher in *C. longiscapa* than in *C. grandiflora* (Fig. S3a).

**Fig.3:**
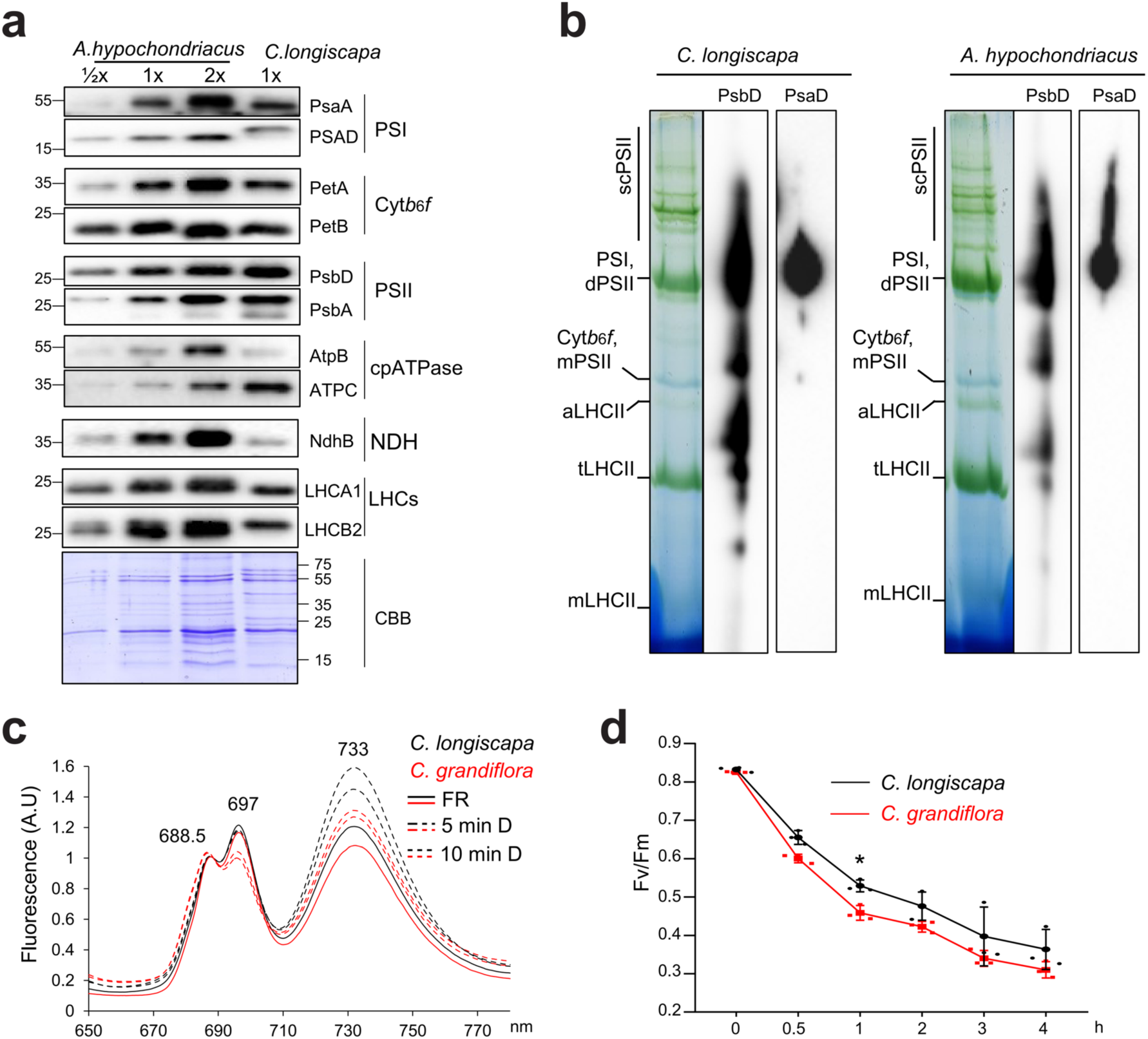
Protein accumulation, complex assembly and acclimation mechanisms of *C. longiscapa*. **(a)** Accumulation of diagnostic protein subunits of the major thylakoidal protein complexes in thylakoids purified from *C. longiscapa* and *A. hypochondriacus* grown under standard light conditions (100 μmol photons m^−2^ s^−1^) for 3 weeks. Samples equivalent to 2 µg of total chlorophyll were loaded for *C. longiscapa* (1×) and resolved in 12.5% PAA gels. Immune detection was conducted by employing antibodies against proteins of the PSII complex (PsbA, PsbD, and LHCB2), the PSI complex (PsaA, PSAD and LCHA1), the Cyt*b*_6_*f* complex (PetA and PetB), the chloroplast ATP synthase (AtpB and AtpC) and the NDH complex (NdhB). As a control for equal loading, a Coomassie-stained PAA gel (CBB) is shown below the immunoblots. The immunoblot experiments were conducted twice (with each replicate corresponding to a pool of 3-5 plants), and similar results were obtained. **(b)** 2D-SDS-PAGE analysis of thylakoids purified from *C. longiscapa* and *A. hypochondriacus* grown under standard light conditions (100 μmol m^−2^ s^−1^) for 3 weeks. Thylakoid samples equivalent to 20 μg of total chlorophyll were solubilized in 1% β-DDM, and resolved in a 6-13.9% gradient gel. SDS-PAGE was performed as second dimension, and immunodetection was conducted with antibodies against PsbD and PSAD as markers for PSII and PSI complexes, respectively. The main photosynthetic complexes are indicated as PSII supercomplexes (scPSII), PSII dimer (dPSII), PSI, PSII monomer (mPSII), Cyt*b*_6_*f*, LHCII assembly (aLHCII), LHCII trimer (tLHCII) and monomeric LHCII (mLHCII). The experiments were performed twice with similar results. **(c)** Chlorophyll a fluorescence spectrum at low temperature (77 °K) in *C. longiscapa* (black) and *C. grandiflora* (red). *C. longiscapa* and *C. grandiflora* grown for 3 weeks under standard conditions were illuminated under far red (FR) light (solid line; state 1) for 2 hours, and then exposed to dark (D) for 5 and 10 minutes, respectively (dotted lines). The emission spectra were recorded at 1 nm s^−1^, with a bandwidth of 5 nm for both excitation and emission. The data were normalized to the peak at 688.5 nm to facilitate comparison of the fluorescence emission peaks derived from PSI. The curves represent the average of 15 curves. n=3 independent biological replicates. **(d)** Analysis of D1 degradation. 3 to 5 detached leaves from *C. longiscapa* and *C. grandiflora* were treated with 1 mM lincomycin. The leaves were exposed to 800 μmol photons m^−2^ s^−1^ for 0, 0.5, 1, 2, 3 and 4 hours. The Fv/Fm was recorded as an indicator of PSII activity. Statistical significance was determined by two-way ANOVA, and *P*-values were adjusted with Sidak’s multiple comparison test, comparing both species at the same time. * *P*<0.05. n=3 independent biological replicates.

Next, we examined the assembly of the main photosynthetic complexes by BN-PAGE in thylakoid samples purified from *C. longiscapa*, *A. hypochondriacus* and *C. grandiflora* (Fig. 3b; Fig. S3b). While comparable accumulation of PSI, PSII dimer (dPSII) and Cyt*b*_6_*f* complexes was observed in all three species, lower accumulation of the intermediate for light-harvesting complex II assembly (aLHCII) was observed in *C. longiscapa*. In addition, a higher accumulation of PSII supercomplexes (scPSII) was observed in *C. longiscapa* compared to *A. hypochondriacus* and *C. grandiflora* (Fig. 3b; Fig. S3b). The higher accumulation of PSII core subunits in *C. longiscapa* prompted us to assess the assembly status of PSII and PSI by two-dimensional gel electrophoresis. 2D-SDS-PAGE analysis revealed a higher accumulation of PSII assembly intermediates and supercomplexes in *C. longiscapa* compared to *A. hypochondriacus* (Fig. 3b). In contrast, *A. hypochondriacus* showed a higher accumulation levels of PSI associated with high molecular weight complexes, most likely due to overaccumulation of PSI-LHCII complexes.

The reversible phosphorylation-dependent movement of light harvesting complex II (LHCII) from PSII to PSI (known as state transitions) and the degradation and replacement of the (photooxidatively damaged) reaction center protein of PSII, PsbA (D1), are important photoprotective mechanisms that fine-tune photosynthesis through energy redistribution between the two photosystems and repair of damaged PSII, respectively (Pribil *et al*., 2018). Due to its very similar leaf morphology (Fig. S4), the sister species *C. grandiflora* was used as a non-extremophile control for the comparative assessment of state transitions and D1 protein turnover. When state transition kinetics were assessed by chlorophyll *a* fluorescence emission at low temperature (77K) (Fig. 3c), similar fluorescence spectra were obtained for *C. longiscapa* and *C. grandiflora* after 2 hours of far-red illumination. The spectra showed the typical peaks for PSII at 688.5 and 697 nm (corresponding to the fluorescence emission of chlorophylls bound to the reaction center proteins CP43 and CP47, respectively) and a peak for PSI at 733 nm (Fig. 3c). At this stage, most of the antennae in both species are associated with PSII. After dark incubation for 5 or 10 minutes, *C. longiscapa* showed faster shifts in the emission peaks than *C. grandiflora* (from 688.5 nm and 697 nm to 733 nm), indicating a faster movement of LCHII from PSI to PSII in *C. longiscapa*. Next, D1 turnover in *C. longiscapa* and *C. grandiflora* was assessed by pre-infiltrating leaves with lincomycin, an inhibitor of plastid translation, and monitoring PSII activity by Imaging-PAM after illumination (400 μmol photons m^−2^ s^−1^). The Fv/Fm value (reflecting the status of functional PSII) decreased more rapidly in *C. grandiflora* than in *C. longiscapa* (Fig. 3d), suggesting greater PSII photodamage in *C. grandiflora* than *C. longiscapa*. Taken together, these results indicate that *C. longiscapa* has higher PSII accumulation levels which may contribute to the better plant acclimation mechanisms compared to *A. hypochondriacus* and *C. grandiflora*.

### *G*enome assembly, gene family expansions, gene duplications and natural variation of *Cistanthe longiscapa*

Since the chromosome content has been widely used as a phylogenetic indicator of plant evolution (Mayrose & Lysak, 2021), we investigated the chromosome content of *C. longiscapa*. Our karyotype analyses showed that *C. longiscapa* has a chromosome number of 2n=22 (Fig. S5a), which is similar to that of *Cistanthe grandiflora* and *Cistanthe ambullatum*, two other species in the Montiaceae family (Marinho *et al*., 2019). Finally, flow cytometry experiments using Arabidopsis as a control allowed us to estimate the genome size of *C. longiscapa*, revealing a value of approximately 539 Mbp (Fig. S5b-d).

Orthologs analysis (BUSCO) (Table S8). The BUSCO analysis of the genome annotation revealed a complete BUSCO of 94.1%, a single BUSCO of 84.9%, and fragmented and missing BUSCOs of 3.2% and 2.7%, respectively (Table S8). Masked repetitive elements in the *C. longiscapa* genome accounted for 44.52% of the genome, with LTRs being the most overrepresented (20.63%) and Gypsy the most abundant (7.44 %) elements (Table S9). Evaluation of the predicted annotated proteins by employing the InterProScan databases revealed that 32,720 predicted proteins were annotated in at least one database, covering 85.74% of the total proteins annotated in the genome (Table S10).

To explore the genomic adaptations that allow *C. longiscapa* to survive in the Atacama Desert, we analyzed the genomic evolution of *C. longiscapa* and compared it with other Caryophyllales and model plant species (Fig. 4). Construction of a phylogenetic tree suggests that *C. longiscapa* diverged from its sister genera approximately 50 million years ago (MYA) (Fig. 4a). KEGG analysis of expanded gene families revealed that, compared to other plant genomes, the *C. longiscapa* genome harbours increased copy numbers of genes associated with DNA repair (including polymerases, endonucleases, and proteins involved in replication), light harvesting and photosynthesis (*plastocyanin 2*, *LHCB1.3*, and *PsbP-like protein 1*, *PPL1*), and autophagy (*ATG1C*, *ATG8D*, *ATG8F*, and *ATG5*) (Fig. 4b; Dataset S1). As gene duplications are known to contribute to plant evolution and environmental adaptation (Panchy *et al*., 2016), we analyzed the GO enrichment of duplicated genes in *C. longiscapa* compared to duplicated genes in other species from the order Caryophyllales and, for comparison, from the model plant Arabidopsis (Fig. 4c). Gene duplications involve 11,384 events in the *C. longiscapa* genome, resulting in the presence of 15,746 duplicated genes. Notably, at least 1,950 of these genes experienced more duplications than the corresponding genes in the common ancestor that gave rise to the Caryophyllales. These duplications were enriched in diverse biological processes including, “organic hydroxy compound metabolic process”, “defense response to salicylic acid”, “response to hypoxia”, “cell death”, as well as “chaperone-mediated protein folding”, “ribosome assembly”, and “stomatal closure” and “photoprotection”. These results suggest that *C. longiscapa* has duplicated genes that allow it to cope with different stresses as well as processes associated with CAM, autophagy, senescence and photosynthesis (Fig.4c; Dataset S1).

**Fig.4:**
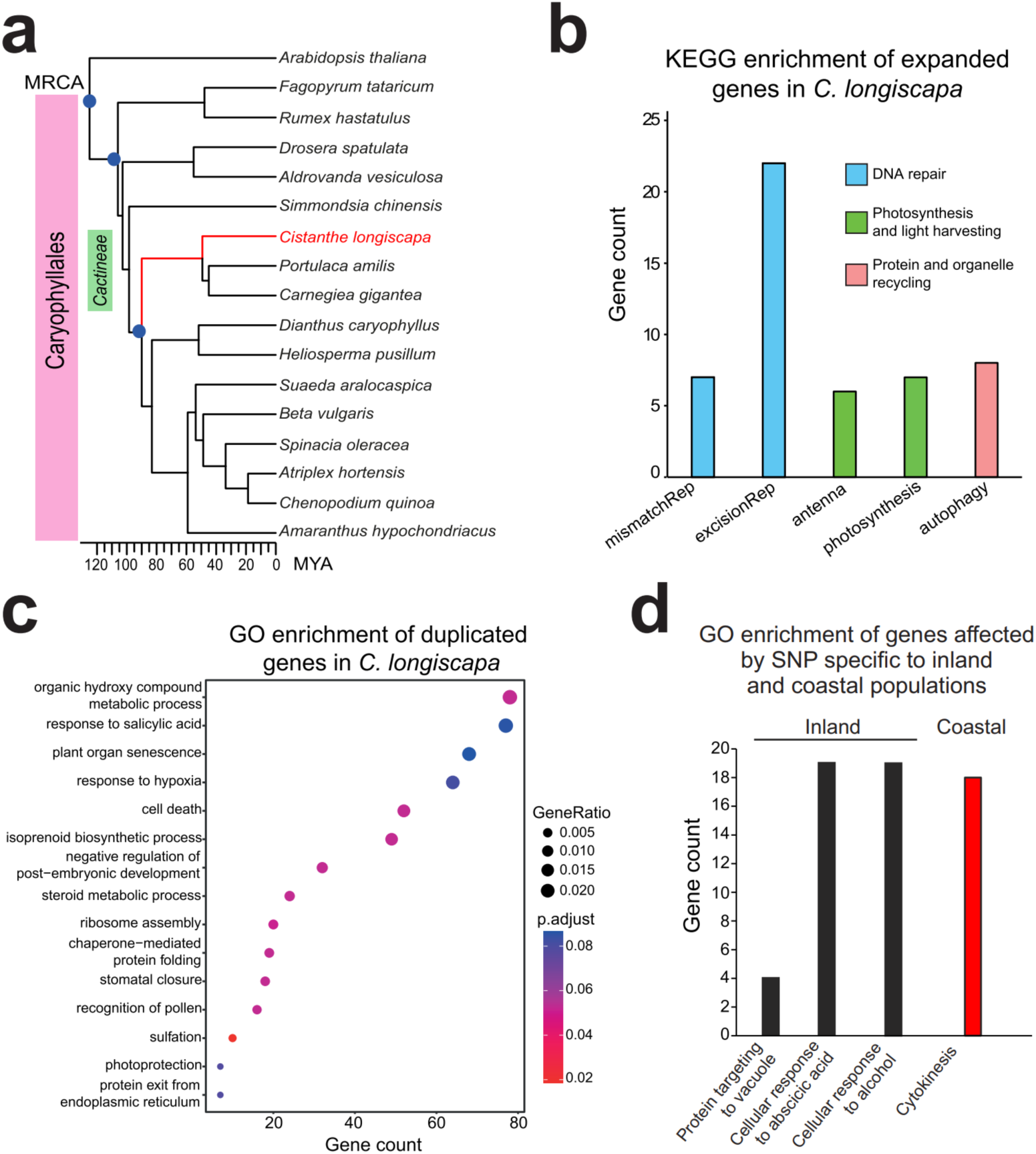
Evolution and genomic adaptations of *Cistanthe longiscapa* in the Atacama Desert. **(a)** Phylogenetic evolution of C. longiscapa compared to other species from the Caryophyllales order. The calibration points are marked in blue. *A. thaliana* represents the outhogroup. MRCA: most recent common ancestor. MYA: million years ago. The information about the genomes of the species analyzed is presented in Table S14 **(b)** Numbers of genes in expanded gene families in the *C. longiscapa* genome. The numbers of additional genes related to the KEGG pathways of photosynthesis, antenna proteins, base excision repair, mismatch repair and autophagy (KEGG codes: map00195, map00196, map03430, map03400, and map04136) are shown. **(c)** GO enrichment analysis of duplicated genes in the *Cistanthe longiscapa* genome. Dot plot shows the GO terms of biological processes identified using EnrichGO. The size of the dots is based on the ratio between gene count and the total number of genes analyzed. The color of the dot indicates the significance of GO enrichment. **(d)** GO terms of biological processes identified using EnrichGO for genes affected by single nucleotide polymorphisms (SNPs) in coastal and inland populations of *C. longiscapa*.

One of the unique features of the Atacama Desert is the climatic variation between inland and coastal regions (Diaz *et al*., 2019). As polymorphisms can be associated with adaptive evolution (Anderson *et al*., 2011), we investigated genomic variation in *C. longiscapa* by whole-genome resequencing of plants collected from the coastal region (location 5) and the inland desert (locations 2 and 3; Table S1), and compared them to the reference draft genome. As expected, large polymorphic variation was observed in samples from the same communities and between coastal and inland populations, due to the self-incompatibility of *C. longiscapa* (Table S11; Table S12) The search for SNPs causing frameshift mutations or premature stop codons revealed that 54 and 18 genes from coastal and inland populations, respectively, were affected (Table S13). The SNPs of putatively affected genes in inland populations showed a single functional GO annotation term that corresponding to cytokinesis, whereas coastal populations showed enrichment of GO terms associated with protein targeting to vacuole, cellular response to abscisic acid stimulus, and cellular response to alcohol (Fig. 4d).

While the plastid genome of *C. longiscapa* had previously been sequenced (Stoll *et al*., 2017b), the structure of the mitochondrial genome was unknown. To complete the genetic information for *C. longiscapa*, we used the Illumina libraries obtained from coastal samples to generate a mitochondrial reference genome (Fig. S6; Dataset S1). The coverage of unique mitochondrial contigs reached a depth of approximately 100x, and a cumulative contig length of ∼410 kpb. The mitochondrial genome of *C. longiscapa* consists of 54 genes, including 30 CDSs, 14 introns, 3 rRNA genes, and 21 tRNA genes. Seed plant mitochondrial genomes typically have multiple genomic configurations, due to the presence of direct and inverted repeats which are recombinationally active (Kozik *et al*., 2019). In *C. longiscapa*, the mitochondrial genome is assembled into three subgenomic circles with sizes of 223,223 bp, 216,604 bp and 135,624 bp for mitochondrial subgenomic configurations 1, 2 and 3, respectively (Fig. S6a-c).

### Transcriptional regulation in the early morning and at midday accounts for most of the diurnal changes in the *C. longiscapa* transcriptome

To investigate the transcriptional responses of *C. longiscapa* to extreme conditions in the Atacama Desert over a 24-hour diurnal cycle, we sampled leaves at four time points every 6 hours from 7:00 am to 1:00 am on the next day (T1 to T4). Sequencing produced an average of 29,552,774 raw reads per sample, with an average mapping rate of 91.1% of the reference genome (Dataset S1). Transcript profiles differed significantly at midday (T2), coinciding with the peak of irradiation in the Atacama Desert (Fig. S7-S8). The most significantly gene expression shifts occurred between midday (T2) and sunrise (T1), with 2,006 up-regulated and 2,050 down-regulated genes, and between sunset (T3) and midday (T2) with 1,090 upregulated and 1,392 downregulated genes (Fig. S8; Dataset S1).

To gain a broader understanding of the genes that underwent up- and down-regulation, we performed an Overrepresentation Gene Ontology analysis of the differentially expressed genes (DEGs) between consecutive time points (Fig.5a). The results revealed a significant number of up-regulated DEGs in the transition from T1 to T2, which are involved in stress-related processes including “response to heat”, “response to high light intensity”, “response to ROS”, and the “ubiquitin-dependent ERAD pathway”. At sunset (T2-T3), one of the most significant GO terms is “regulation of stomatal movement”, in line with the property of CAM plants to open their stomata at night. As expected, genes associated with “response to heat” were down-regulated at sunset. Finally, the GO terms “cellular response to abiotic stimulus”, “cellular response to environmental stimulus”, “cellular response to hypoxia” and “response to red and far-red light” were up-regulated at midnight and dawn (Fig. 5a).

**Fig. 5:**
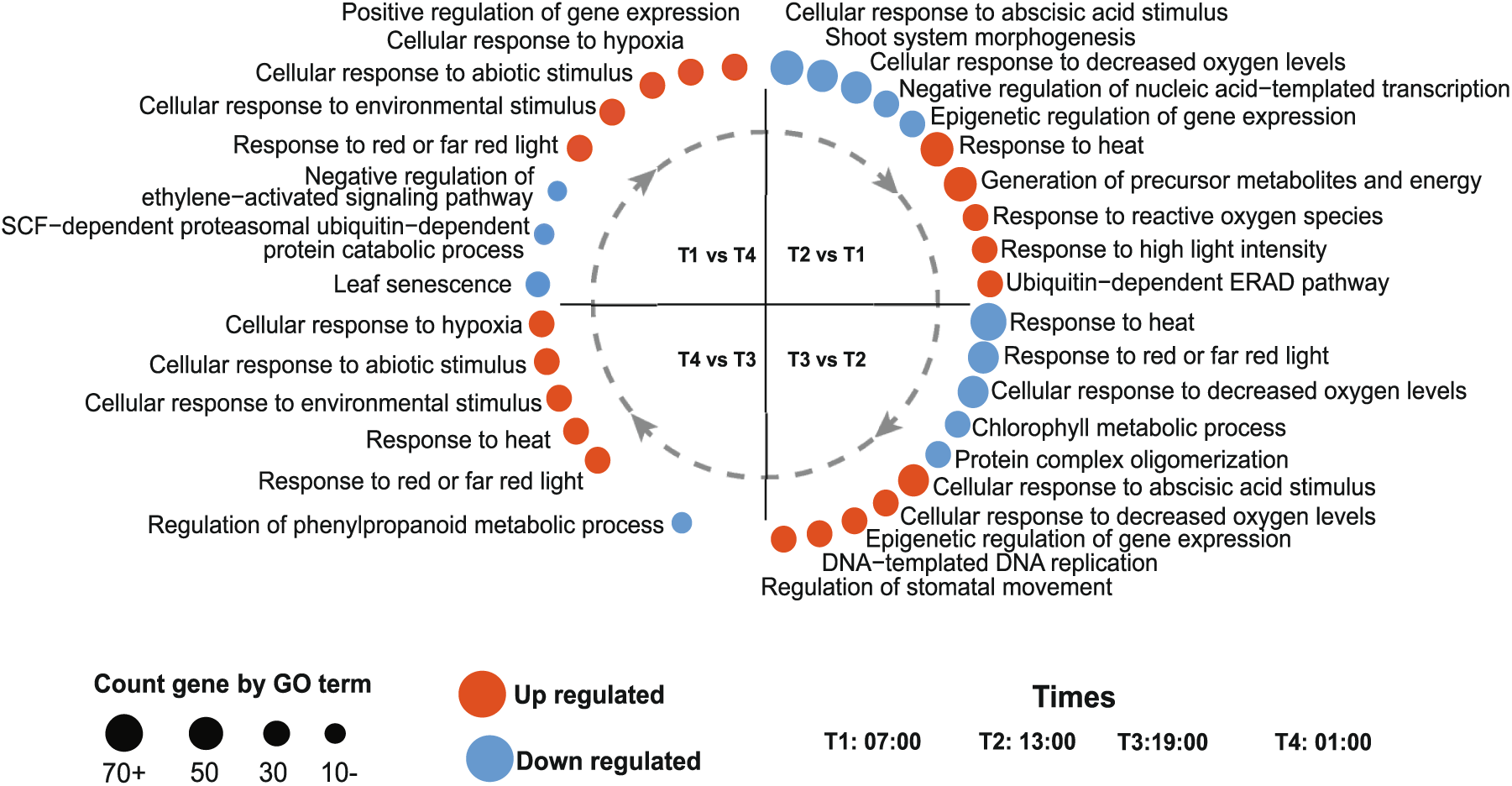
Time-resolved transcriptomic of *C. longiscapa*. Enrichment of gene ontology (GO) terms for specific biological processes at each analyzed time point. Overrepresented terms from up- and down-regulated DEGs are shown as red and blue dots, respectively. The dot size represents the gene count for the enriched term. All GO terms shown have an overrepresentation with an adjusted *P*-value < 0.05.

Transcriptional regulation of genes associated with heat and drought stress, nutrient limitation and high UV irradiance are some of the conserved responses among plants living in the Atacama Desert (Eshel *et al*., 2021). Therefore, we analyzed the accumulation of transcripts involved in these processes in our time-resolved datasets. As expected, we observed a strong up-regulation of transcripts encoding for heat shock proteins (e.g. *HSP17* and *HSP90*), and drought response (*RD19A*, *ROF2* and *MBF1C*) in T2 compared to T1 (Dataset S1). Similarly, genes associated with nitrogen and sulphur use (e.g. *NPF8.1*, *GLN1;4* and *SDI1*) and UV responses (e.g. *UVR8* and the wax-related *CER1)* were also up-regulated at midday (Dataset S1). In contrast, transcripts of hormone-related genes (e.g. *LOG5* and *IAGLU*) and cell wall biosynthesis-related genes (e.g. *ATCSLG3*, *CSL3*, and *IRX15)* were up-regulated during the afternoon and night (Dataset S1). These results suggest that the daytime responses of *C. longiscapa* are related to stress resistance, whereas the nighttime responses promote plant growth.

### *Cistanthe longiscapa* exhibits CAM metabolism in the Atacama Desert

CAM photosynthesis has been studied for decades, because this pathways increases water use under extreme drought conditions. Therefore, we analyzed the accumulation of transcripts encoding for CAM-related proteins in *C. longiscapa*. Total acid titration (Fig. 6a) and malate content (Fig.6b) of plants harvested in the Atacama Desert showed the expected pattern reported for CAM species, with a peak in malate at sunrise (T1), a decrease during the day (T2 and T3) and a subsequent increase at night (T4). In addition, the transcript encoding CAM-related enzymes, including carbonic anhydrase (*CA*), phosphoenolpyruvate carboxylase (*PPC*) and malate dehydrogenase (*MDH*), showed high accumulation at the beginning of the day (T1) and during the night (T4) (Fig. 6b). In contrast, transcripts encoding enzymes involved in decarboxylation, including NADP-malic enzyme (*NADP-ME*) and phosphoenolpyruvate carboxykinase 1 (*PPCK*) were up-regulated during the day (Fig. 6b). These results indicate that *C. longiscapa* performs CAM photosynthesis in the Atacama Desert.

**Fig.6:**
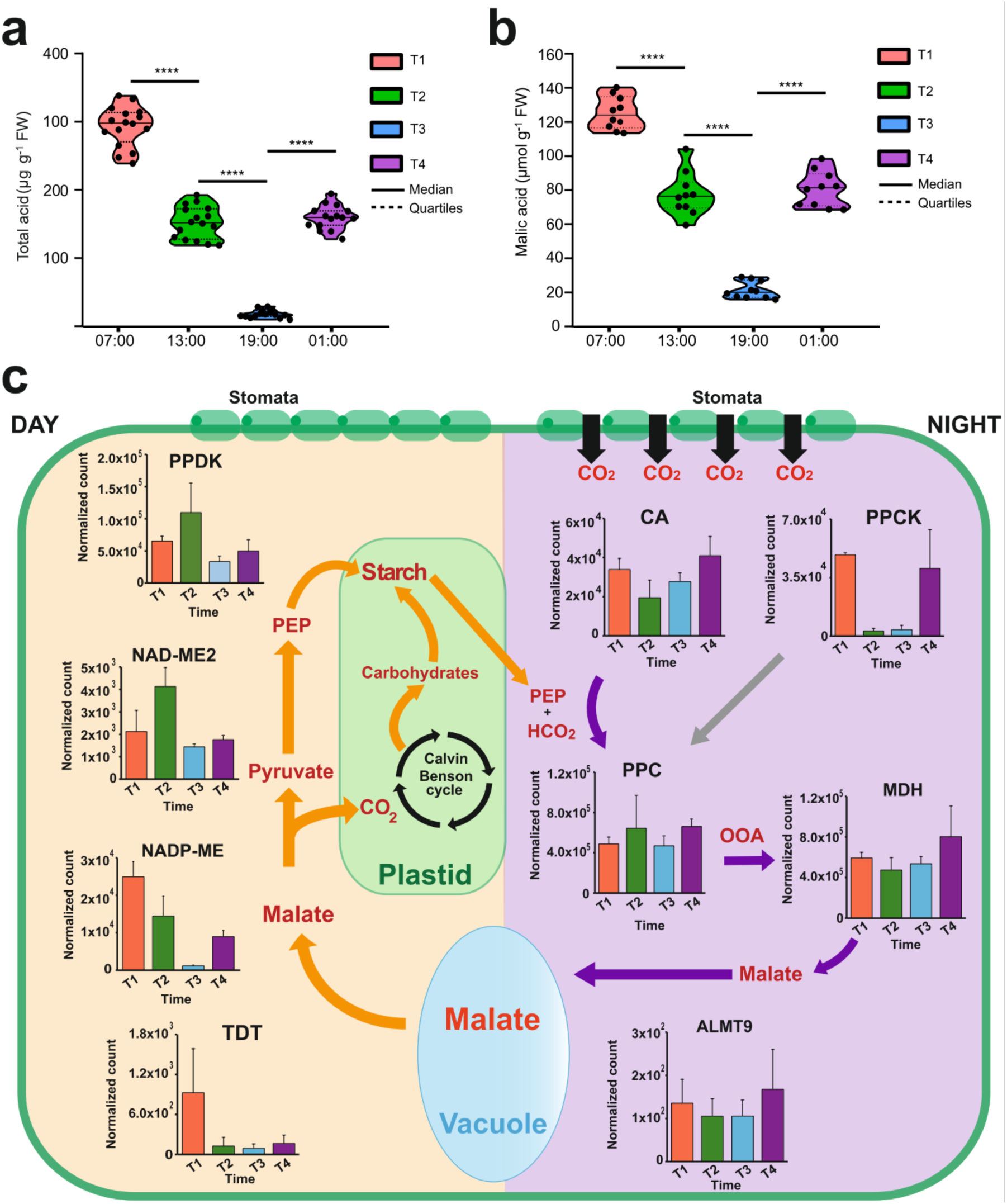
C. *longiscapa* performs CAM photosynthesis in the Atacama Desert. Quantification **a**) total acid and **b**) malic acid contents of leaves of *C. longiscapa* plants grown in the Atacama Desert at four different timepoints were measured. The leaves were harvested at sunrise (7:00; T1), midday (13:00; T2), sunset (19:00; T3) and night (1:00; T4). The violin plot indicates the median (solid lines) and the quartiles (dotted lines) for each group of samples. The total acid and malic acid contents were normalized to fresh weight (FW). Statistical significance was tested by one-way ANOVA, and P-values were adjusted with Sidak’s multiple comparison test. **** P<0.0001 (n=10 biological replicates). **c**) Simplified view of the CAM photosynthetic pathway. Text in red color indicates key metabolites involved in the CAM pathway, grey arrows represent alternative routes, purple arrows indicate pathway steps occurring predominantly during low-light hours, and orange arrows indicate pathway steps occurring predominantly during daylight hours. The bar plots show DESeq2’s median of ratios normalized counts for each timepoint sampled for crucial enzymes and transporters related to CAM. The plots include the normalized transcript abundance for the carbonic anhydrase (*CA*), pyruvate orthophosphate (Pi) dikinase (*PPDK*), phosphoenolpyruvate carboxylase (*PPC*), malate dehydrogenase (*MDH*), aluminum-activated malate transporter 9 (*ALMT9*), tonoplast dicarboxylate transporter (*TDT*), NADP-dependent malic enzyme (*NADP-ME*), NAD-dependent malic enzyme 2 (*NAD-ME2*), and phosphoenolpyruvate carboxykinase (*PPCK*).

### Transcripts coding for plastid localized proteins are largely regulated in the Atacama Desert

We explored the differences in transcript accumulation by clustering the DEG transcripts according to similar expression patterns (Fig. 7b). Of the eight clusters, two clusters peaked at sunrise and four clusters showed gene expression peaks at midday (Fig. 7b). While none of the clusters showed transcript peaks at midnight, only one cluster showed increased gene expression at sunset. The largest transcriptional changes were observed at midday, and included clusters 3 (25 genes), 4 (160 genes), 5 (144 genes), and 6 (1,024 genes). In particular, the clusters showed a high accumulation of transcripts encoding plastid-localized proteins. Therefore, we examined the differences in transcript accumulation by similar expression patterns and classified them by function (Fig.7b; Dataset S1).

**Fig. 7:**
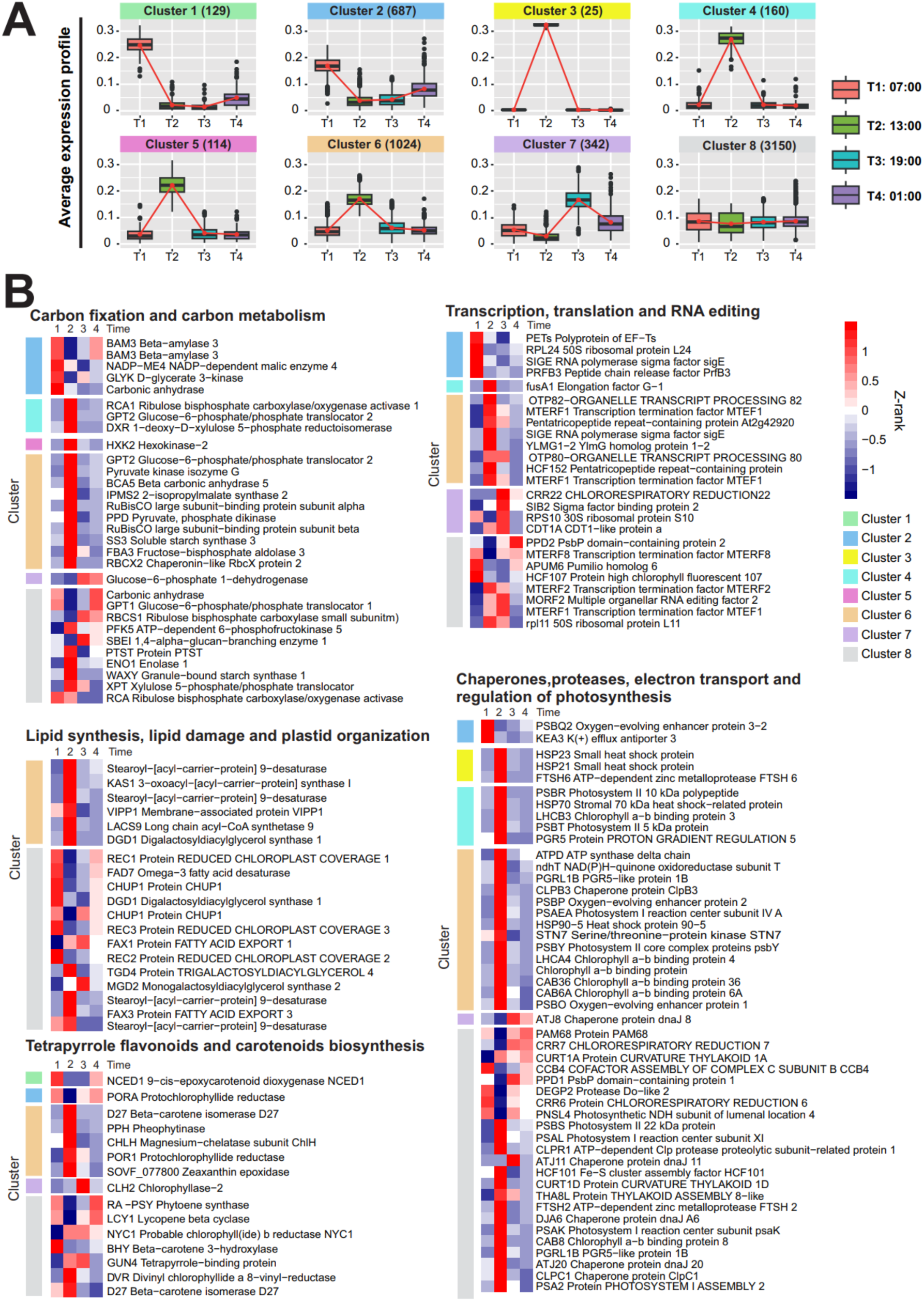
Differentially expressed transcripts of *Cistanthe longiscapa*. **(a)** Clusters of differentially expressed genes for the four time points. Differentially expressed genes were clustered based on their average expression profiles. The clusters were generated using the k-means algorithm with a value of k=8 and 1000 bootstrap terations. The input data for clustering were derived from logarithmically transformed counts for each time point, including their respective replicates. **(b)** Heat map showing the expression of differentially expressed transcripts coding for chloroplast localized-proteins. The MAKER annotation was employed to select transcripts coding for chloroplast-localized proteins, and were grouped by biological function: Carbon fixation and carbon metabolism; Lipid synthesis, lipid damage and plastid organization; Tetrapyrrole, flavonoids and carotenoids biosynthesis; Transcription, translation and RNA editing; Chaperones, proteases, electron transport and regulation of photosynthesis. The hierarchical clustering was generated using the hclust complete linkage method in z-scored “DESeq2’s median of ratios” mean expression values. Low expression is indicated in blue, high expression in red. The cluster to which the transcripts belong are indicated in the left.

For carbon metabolism, the transcripts that accumulate upon light onset largely correspond to proteins known to appear early during daytime (*BAM1*, *NADP-ME4*). While DEGs related to carbon fixation (*FBA3, DXR* and *HXK2*), starch accumulation (*SS3*), Rubisco activase (*RCA*) and Rubisco assembly (*RBCX2*) were up-regulated at midday, slight changes in transcripts encoding starch granule bound synthase (*WAXY*) and starch branching enzyme (*SBEI*) were observed at midday and noon, respectively (Fig 7b). The transcripts encoding for lipid biosynthetic enzymes (*KAS1*, *LACS9* and *DGD1*) accumulate largely at midday (Fig.7b). Similarly, the large midday accumulation of the stress-induced transcript encoding for *VIPP1* may indicate active lipid repair during high light periods in the Atacama Desert (Gupta et al., 2021). While *VIPP1* transcript accumulation decreases towards the end of the day, we observed up-regulation of lipid biosynthetic transcripts such as *MDG2* (Murakawa et al., 2019), consistent with MGDG biosynthesis under low light (Yu *et al*., 2021). Interestingly, many of the transcripts encoding proteins involved in plastid morphology (*CHUP1*) and plastid movement (*REC3*) showed similar expression patterns to transcripts involved in lipid biosynthesis (Fig.7b), suggesting a close link between these processes (Larkin et al., 2016; Oikawa et al., 2008). Upregulation for the *NCED1* (cluster 1) and *Zeaxanthin epoxidase* (cluster 6) transcripts, may indicate active ABA biosynthesis under early light and at midday, consistent with responses under high light, drought and heat stress (Tan et al., 1997). Transcripts associated with chlorophyll biosynthesis (*CHLH* and *POR1*; cluster 6) and degradation (*CLH2*; cluster 7) peaked during the day and at midday, which may indicate active repair mechanisms of chlorophyll-containing complexes (e.g. PSII and PSI) (Wang and Grimm, were slightly up-regulated during low light periods (T1 and T4), and followed a similar pattern to that observed for transcripts involved in lipid biosynthesis (Fig. 7b), which has been tentatively attributed to the lipophilic nature of these compounds (Torres-Montilla and Rodriguez-Concepcion, 2021).

The DEGs that accumulate at the onset of light largely correspond to transcripts encoding ribosomal proteins (*RPL24*), plastid-localized sigma factors (*SIGE*), and translation elongation (*PETs*) and mRNA processing (*PRFB3*). However, the accumulation of most of these transcripts decreased towards the night, which is attributed to the circadian regulation of genes encoding plastid-localized proteins involved in transcription and translation (Noordally *et al*., 2013).

Finally, we examined the accumulation of transcripts encoding proteins involved in protein folding, degradation, electron transport and regulation of photosynthesis (Fig. 7b). The transcripts encoding *KEA3* peaked early upon illumination and may contribute to non-photochemical quenching mechanisms that occur at midday (Uflewski et al., 2024). Cluster 3 contained genes related to protein folding and degradation, including the transcripts encoding chloroplast-localized *HSP23* and *HSP21*, and a member of a chloroplast ATP-dependent zinc metalloprotease family (*FTSH6*). Similarly, transcripts for chlorophyll-binding proteins (*LHCB3*, *LHCA4*, *CAB6* and *CAB36*) were up-regulated at midday (Fig. 7b). The high expression of *PGR5* (cluster 4) coincided with the expression of chaperones and proteases at midday, consistent with the proposed role of PGR5 in photoprotection (Rantala *et al*., 2020). In addition, transcripts encoding for *STN7* (cluster 6) and *CURT1A* (cluster 8) were up-regulated at midday, supporting photosynthetic acclimation mechanisms at this time (Tikkanen *et al*., 2012; Pribil et al., 2018). Finally, the expression of PSII and PSI assembly factors at dusk (*PAM68* and *PPD1*, respectively), and the accumulation of transcripts encoding proteases at low light levels (*DEG2*) are tentatively attributed to the repair of damaged PSII and PSI accumulated during the day (Fig. 7b).

Overall, these results highlight the central role of chloroplast-associated processes in regulating the resilience of *C. longiscapa* in the Atacama Desert.

## Discussion

We have explored the physiological, genomic and transcriptomic responses that contribute to the survival of *C. longiscapa* under the Atacama Desert. Our photosynthetic measurements showed a better Φ_II_ in *C. longiscapa* compared to *C. grandiflora* and *A. hypochondriacus*, which may be due to the higher content of assembly intermediates and PSII complexes associated in super complexes reducing the photoinhibition of PSII at high light intensities (Che *et al*., 2020). Similarly, the higher content of PSII may contribute to the photoprotection during plant adaptation to drought and high light (Tikkanen *et al*., 2014). Consistent with the expansion of genes encoding antenna proteins (*LHCB1.3*) in the genome and the high expression levels of LHCII transcripts at midday, we also observed faster antenna movements in *C. longiscapa*, supporting the role of state transitions under challenging environmental conditions to rebalance the energy distribution between photosystems (Pribil *et al*., 2018). Consistent with this hypothesis, our transcriptome analyses also revealed high accumulation levels of the *STN7 kinase* mRNA (responsible for LHCII phosphorylation (Tikkanen *et al*., 2012)) at midday. Together with the increased ability to perform state transitions, the expansion of the plastocyanin gene family may also contribute to reducing the excitation pressure on PSII, by enhancing linear electron transfer to PSI (Schottler *et al*., 2004). Similarly, the high level of PGR5, and the duplication of *PGRLB1* in the *C. longiscapa* genome are likely to contribute to enhanced photoprotection (Rantala *et al*., 2020). The expansion of the PsbP-like protein family, which has been linked to PSII biogenesis and the accumulation of PSII supercomplexes and their optimized performance under fluctuating light conditions (Che *et al*., 2020) is consistent with the observed high accumulation of PSII intermediates and supercomplexes, which may provide a larger pool of PSII in *C. longiscapa* to avoid PSII photoinhibition and allow for faster state transitions (Dietzel *et al*., 2011; Kim *et al*., 2020). This interpretation is further supported by our analysis of D1 protein degradation.

We observed a large expansion of gene families related to DNA repair, photosynthesis, and autophagy. Previous work in maize has shown that autophagy plays a critical role in plant lipid and secondary metabolism (McLoughlin *et al*., 2018). Furthermore, overexpression of *ATG18a* in apple has been shown to increase thermotolerance, enhance photosynthesis, and activate genes involved in ROS detoxification and drought stress at high temperatures (Huo *et al*., 2020). Therefore, the expansion of autophagy-related gene families in *C. longiscapa* (e.g., *ATG8*, *ATG1*, *ATG5*, and *ATG18a*, which is present in three copies in the genome; Dataset S1), may indicate that autophagy is one of the major evolutionary mechanisms facilitating plant adaptation to the harsh environmental conditions of the Atacama Desert. Similarly, enhancing DNA repair capacity in the *C. longiscapa* genome (e.g., by increasing the copies of the *FEN1* and *POLD* genes; Dataset S1) may contribute to maintaining DNA integrity under high irradiance and high ROS accumulation. A similar mechanism has previously been proposed for a Desert green alga, *Chlorella spec.*, where adaptations in photosynthesis and DNA repair are largely responsible for the extraordinary UV tolerance (Wang *et al*., 2022). Similarly, the ROS-scavenging properties of betalains (Sadowska-Bartosz & Bartosz, 2021) which largely accumulate in the epidermal cell layer of *C. longiscapa* may also contribute to minimizing oxidative stress (Polturak *et al*., 2017), and act as an additional mechanism of photoprotection. In addition, the distinct SNP profiles in coastal and inland plant populations suggest substantial genetic adaptation to the local environment. Evidence for such genetic adaptation is found, for example, in genes involved in cytokinesis and phenylpropanoid biosynthesis. In contrast, the SNP profiles of inland plants suggest adaptation through enhanced abiotic stress tolerance and metabolic flexibility, likely shaped by the more stable but stressful and resource-poor inland environment (Diaz *et al*., 2019). Together, these genomic changes are likely to contribute to enhanced plant tolerance to adverse environmental conditions in the desert (Izumi *et al*., 2017; Eshel *et al*., 2022).

The high nocturnal malate levels overlapped with the up-regulation of genes involved in CO₂ fixation and storage, such as phosphoenolpyruvate carboxylase (*PPC*), malate dehydrogenase (*MDH*) and tonoplast dicarboxylate transporters (*TDT*). In contrast, the reduction in malate levels during the day reflects decarboxylation processes that release CO₂ for photosynthesis (Winter & Smith, 2022). This is supported by the up-regulation of NADP-malic enzyme (*NADP-ME*) and phosphoenolpyruvate carboxykinase (*PPCK*), which enable CO_2_ fixation by Rubisco (Winter & Smith, 2022). Similarly, transcripts encoding enzymes of the CBB cycle, including hexokinase (*HXK2*) and fructose−bisphosphate aldolase (*FBA3*), and starch synthesis (e.g. soluble starch synthase 3, *SS3*) are up-regulated at midday. In addition, the high transcript accumulation and gene duplication of Rubisco activases (three copies in the genome; Dataset S1) may contribute to photosynthesis in the Atacama Desert (Waheeda *et al*., 2023). These oscillations in malate levels and transcript accumulation confirmed CAM photosynthesis in *C. longiscapa* in the Atacama Desert (Holtum *et al*., 2021).

In summary, our work highlights key mechanisms regulating photosynthesis and plant Desert. Future research will focus on characterizing the mechanisms of carbon utilization and light acclimation in *C. longiscapa* by generating mutants to validate the roles of the genes identified in this study. This work advances our understanding of chloroplast development and photosynthesis, contributing to fundamental science and agricultural strategies to enhance plant resilience under extreme environmental conditions.

**Table 1.** Genomic features of *Cistanthe longiscapa*. The table shows major genomic features of *C. longiscapa* in comparison to the sequenced genomes of the model plant *Arabidopsis thaliana* and other members of the order Caryophyllales: *Portulaca amilis*, *Amaranthis hypochondriacus*, *Beta vulgaris*, *Chenopodium quinoa* and *Simmondsia chinensis*.

| Feature | *Cistanthe longiscapa* | *Arabidopsis thaliana* | *Portulaca amilis* | *Amaranthus hypochondriacus* | *Beta vulgaris* | *Chenopodium quinoa* | *Simmondsia chinensis* |
| --- | --- | --- | --- | --- | --- | --- | --- |
| **Genome version** | V1.0 | Araport11 | V1.0 | V2.1 | EL10_1.0 | V1.0 | V1.0 |
| **Genome size (Mbp)** | 530 | 135 | 403.89 | 466 | 540 | 1385 | 886.73 |
| **Ploidy** | diploid | diploid | diploid | diploid | diploid | allotetraploid | tetraploid |
| **Chromosomes** | n = 11 | n = 5 | n = 9 | n = 16 | n = 9 | n = 18 | n = 26 |
| **GC (%)** | 38.39 | 36 | 37.05 | 33.4 | 36.14 | 36.91 | 37.28 |
| **Protein coding genes** | 38,153 | 27,416 | 52,037 | 23,847 | 24,255 | 53,436 | 23,490 |
| **Average gene length (bp)** | 4,121 | 3,373 | 3,071 | 4,858 | 6,139 | 4,797 | 7,864 |
| **Genome covered with genes (%)** | 29.65 | 54.85 | 39.57 | 28.7 | 28.73 | 15.51 | 20.83 |
| **Duplicated genes** | 15,746 | 10,131 | 30,710 | 8,530 | 11,158 | 42,337 | 12,956 |
| **Duplicated genes (%)** | 41.27 | 36.95 | 59.02 | 35.77 | 46.00 | 79.23 | 55.16 |
| **Genome covered with TE (%)** | 44.52 | 19.9 | 46.96 | 48 | 42.3 | 64 | 65.73 |
| **Photosynthetic metabolism** | CAM | C3 | C4/CAM | C4 | C3 | C3 | C3 |
| **Self-fertilization** | no | yes | yes | yes | no | yes | no |
| **Succulent leaves** | yes | no | yes | no | no | no | yes |
| **Lifecycle** | annual | annual | annual | annual | biennial | annual | evergreen |
| **Complete BUSCOs (%)\*** | 94.1 | 98 | 89.8 | 80.2 | 92.8 | 95 | 88.7 |
| **Fragmented BUSCOs (%)** | 3.2 | 1.2 | 6.9 | 8.7 | 2.6 | 2.7 | 7.8 |
| **Missing BUSCOs (%)** | 2.7 | 0.8 | 3.3 | 11.1 | 4.6 | 2.3 | 3.5 |

\* BUSCO values are from the complete set of annotated proteins for each genome according to the database Embryophyta\_odb10.

## Acknowledgments

We thank Ivonne Feldman, Reinaldo Campos and Julian Verdonk for collecting seeds and plant samples; OSI and RB thank Karin Köhl and Stephanie Ruf for advice on plant cultivation and seed propagation. This research was financially supported by ANID – Millennium Science Initiative Program - ICN2021_044 (to AO) and the Max Planck Society (to RB).

## Author Contributions

OSI, CM, RB and AO designed the experiments. PTR, AR. and CM performed the genome assembly and annotation, SC performed the mitochondrial genome assembly and annotation. RY extracted nuclei and performed flow cytometry analyses. CB performed the karyotype analyses. PO, RNP, DO and AAM collected and processed samples for RNA and DNA library construction, and malate analyses. AM performed the HPLC and analyzed the data. OSI analyzed the photosynthetic performance, conducted the immunoblots, BN-PAGE, 2D-SDS-PAGE, the analysis of state transitions and the analysis of D1 degradation. RY, AAM, OSI, and AO obtained the photographs. AAM, RNP, AZS, AM, FBH, ADG, MLA, MG, AM, MM, and RAG provided critical experimental suggestions and feedback on the draft manuscript. PTR, OSI, AR, CM, RB and AO drafted the manuscript. CM, RB and AO secured the funding. All authors commented on and approved the manuscript.

## Data availability

All the data are included in the article and/or SI Appendix. Plant material is available on request from the corresponding authors. The SRA and raw data have been deposited at NCBI BioProject ID PRJNA1075442. The Whole Genome Shotgun (WGS) project which includes the nuclear and mitochondrial genomes have been deposited at DDBJ/ENA/GenBank under the accession JBAIVK000000000. The version described in this paper is the JBAIVK010000000 version.

---

**Fig. S1:**
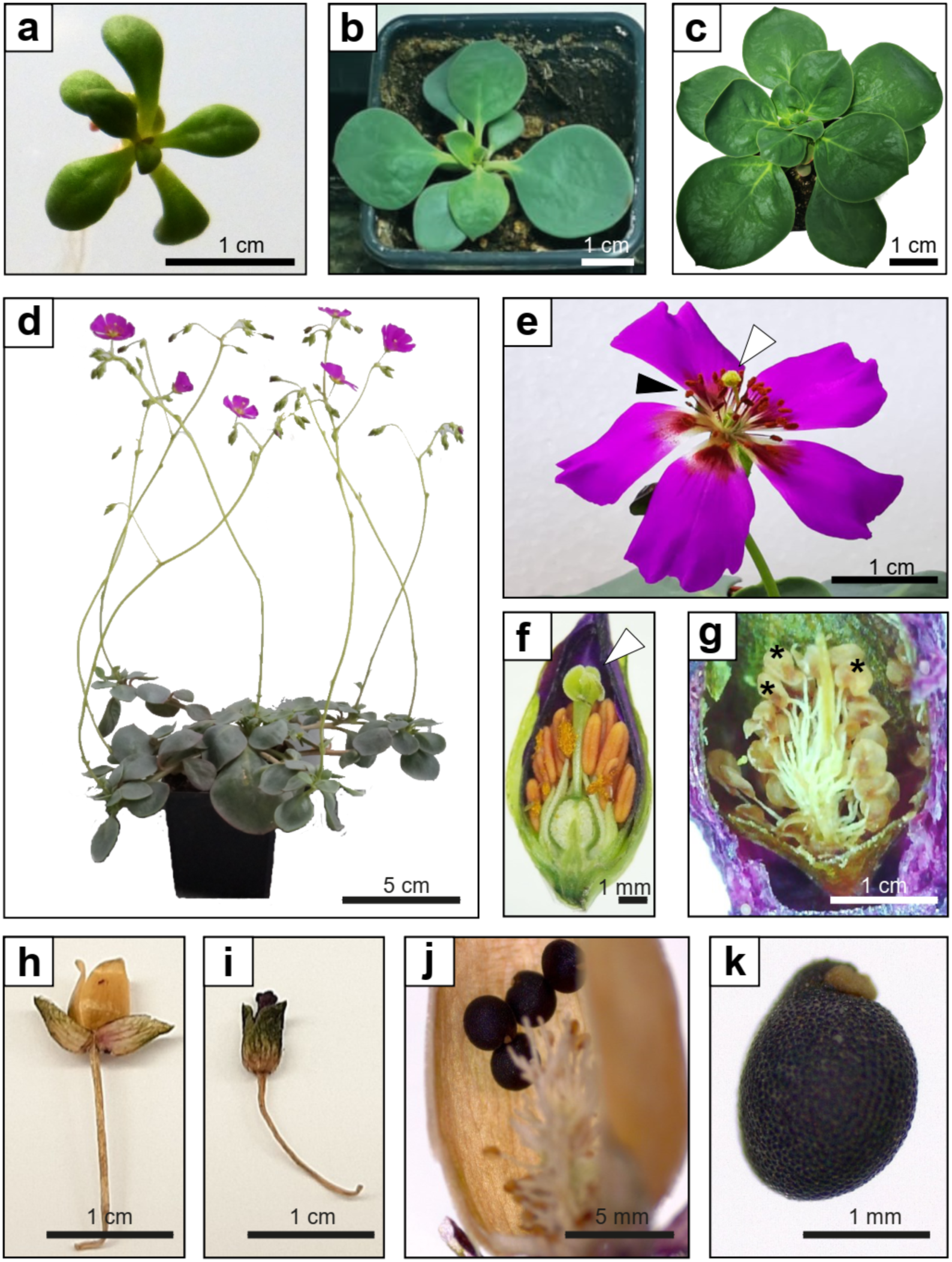
Morphological features and developmental stages of *Cistanthe longiscapa*. Vegetative phase of *C. longiscapa*. **a**) 3 weeks after germination, **b**) 6 weeks after germination and **c**) 10 weeks after germination. **d**) Reproductive phase of *C. longiscapa* 15 weeks after germination. In the life cycle of C. longiscapa, about 50 flowers can be produced. **e-f**) Floral bud and flower of *C. longiscapa*. Anthers and stigma are indicated with black and white arrows, respectively. **g**) Embryo development after fertilization. Three embryos are indicated by black asterisks. **h**) Fertilized capsule of C. longiscapa. **i**) Aborted (non-fertilized) flower of C. longiscapa. The capsule does not form, and flower abortion occurs at the base of the pedicel. **j**) Seeds inside the capsule. About 120 seeds can be collected from one flower. **k**) Seed of *C. longiscapa*.

**Fig. S2:**
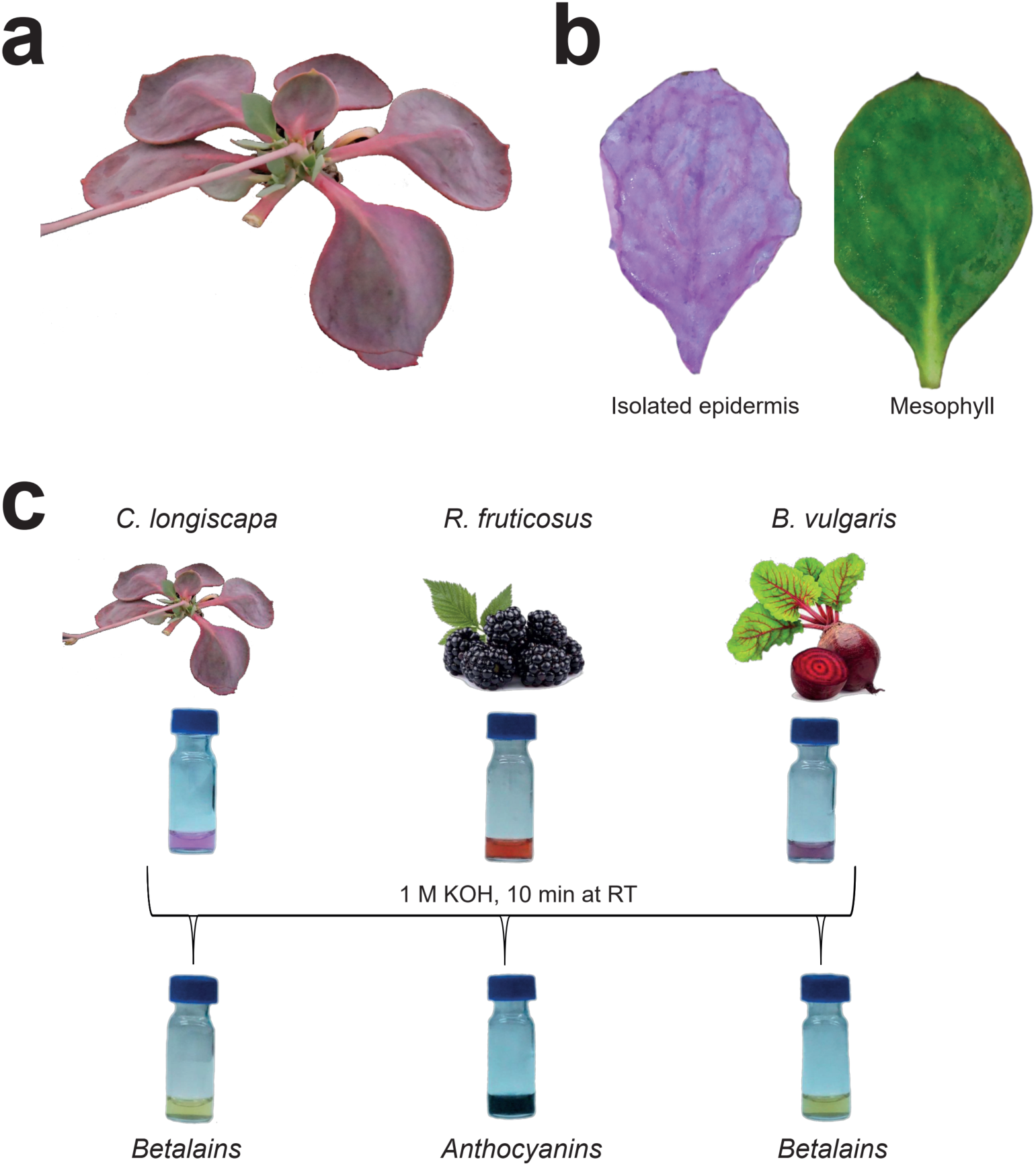
Drought-induced accumulation of betalains in the epidermal layer of *Cistanthe longiscapa* leaves. **a)** Accumulation of pigments in leaf tissue of *C. longiscapa* upon water-deprived growth. Plants were grown for 12 weeks under standard conditions and then water was withheld for 2 weeks. **b)** Epidermal layer (left) and leaves after removal of the epidermis (right) of *C. longiscapa* grown under the conditions described in a). Note that the pigments accumulated exclusively in the epidermal layer. **c)** Biochemical assessment of pigment composition. Water-soluble pigments were extracted from *C. longiscapa*, *Beta vulgaris* (*B. vulgaris*) and *Rubus fruticosus* (*R. fruticosus*) (see Method section). The colorimetric assay indicates the presence of betalains (yellow) or anthocyanins (dark purple) in the vials.

**Fig. S3:**
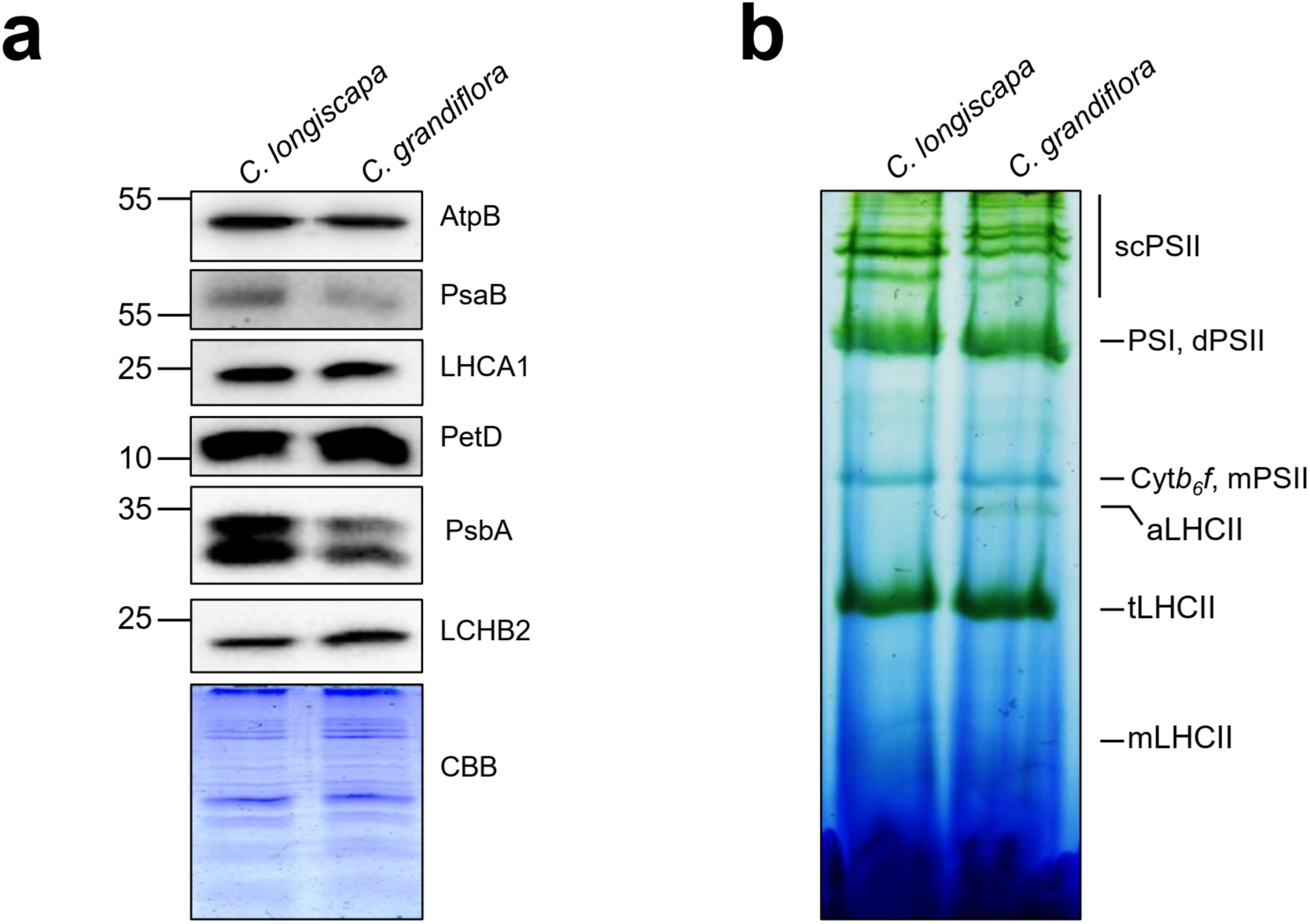
Protein accumulation and complex assembly in thylakoid samples from *Cistanthe longiscapa* and *Cistanthe grandiflora*. **a)** Protein accumulation of diagnostic protein subunits of the major thylakoidal protein complexes in thylakoids purified from *C. longiscapa* and *C. grandiflora* grown under standard light (100 µmol m^−2^ s^−1^) for 3 weeks. Samples equivalent to 2 µg of total chlorophyll were loaded and resolved by electrophoresis in 12.5% PAA gels. Immunochemical detection was conducted by employing antibodies against proteins of the PSII complex (PsbA and LHCB2), the PSI complex (PsaB and LCHA1), the Cyt*b*_6_*f* complex (PetD), the chloroplast ATP synthase (AtpB). As a control for equal loading, a Coomassie-stained PAA gel (CBB) is shown below the series of immunoblots (n = 2 biological replicates, with each replicate corresponding to a pool of 3-5 plants). **b)** BN-PAGE analyses of thylakoids purified from *C. longiscapa* and *C. grandiflora* grown under standard light (100 μmol photons m^−2^ s^−1^) for 3 weeks. Thylakoid samples equivalent to 10 µg of total chlorophyll were solubilized in 1% β-DDM, and resolved in an 6-13.9% gradient gel. The main photosynthetic complexes are indicated as PSII supercomplexes (scPSII), PSII dimer (dPSII), PSI, PSII monomer (mPSII), Cyt*b*_6_*f*, LHCII assembly (aLHCII), LHCII trimer (tLHCII) and monomeric LHCII (mLHCII). n=2 independent biological replicates.

**Fig. S4:**
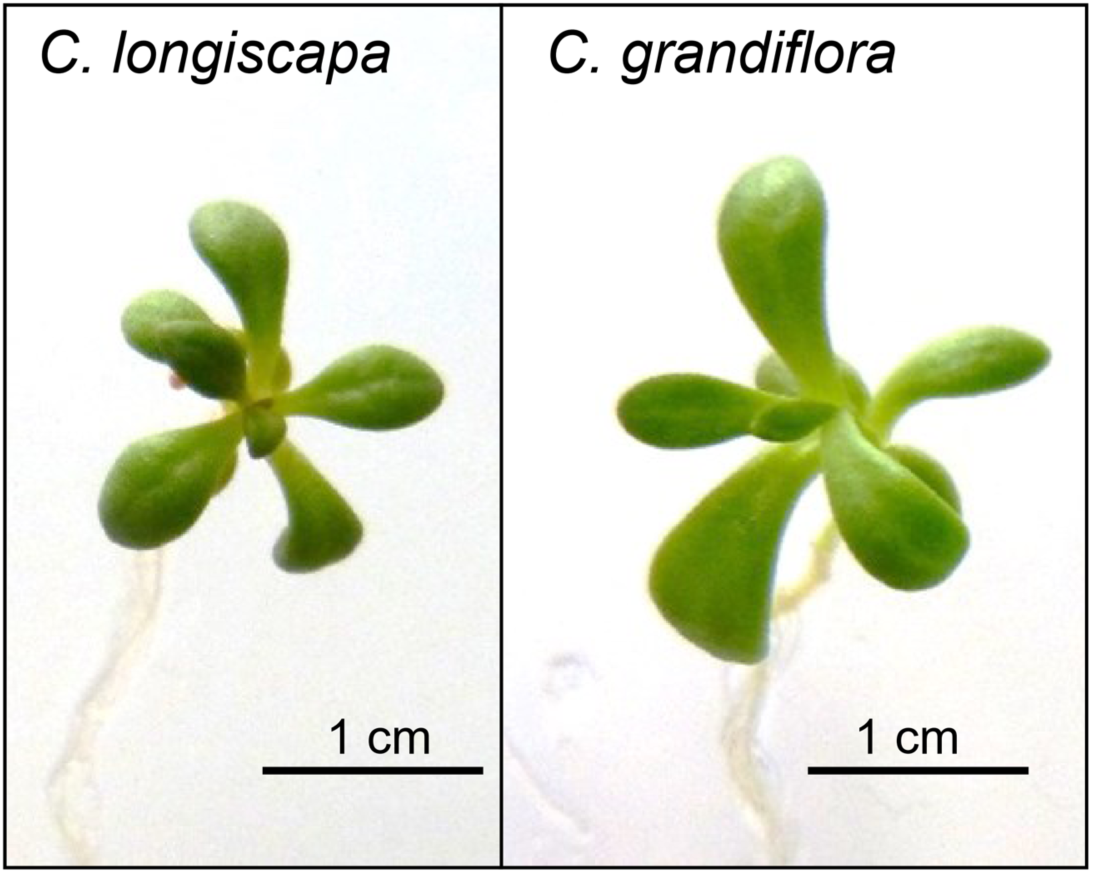
Leaf morphology and plant sizes of *Cistanthe longiscapa* and *Cistanthe grandiflora*. Representative pictures of plant morphology of *Cistanthe longiscapa* and *Cistanthe grandiflora* after 3 weeks of cultivation under standard *in vitro* conditions (see Methods). Note similar plant size and similar leave shape of the two species.

**Fig. S5:**
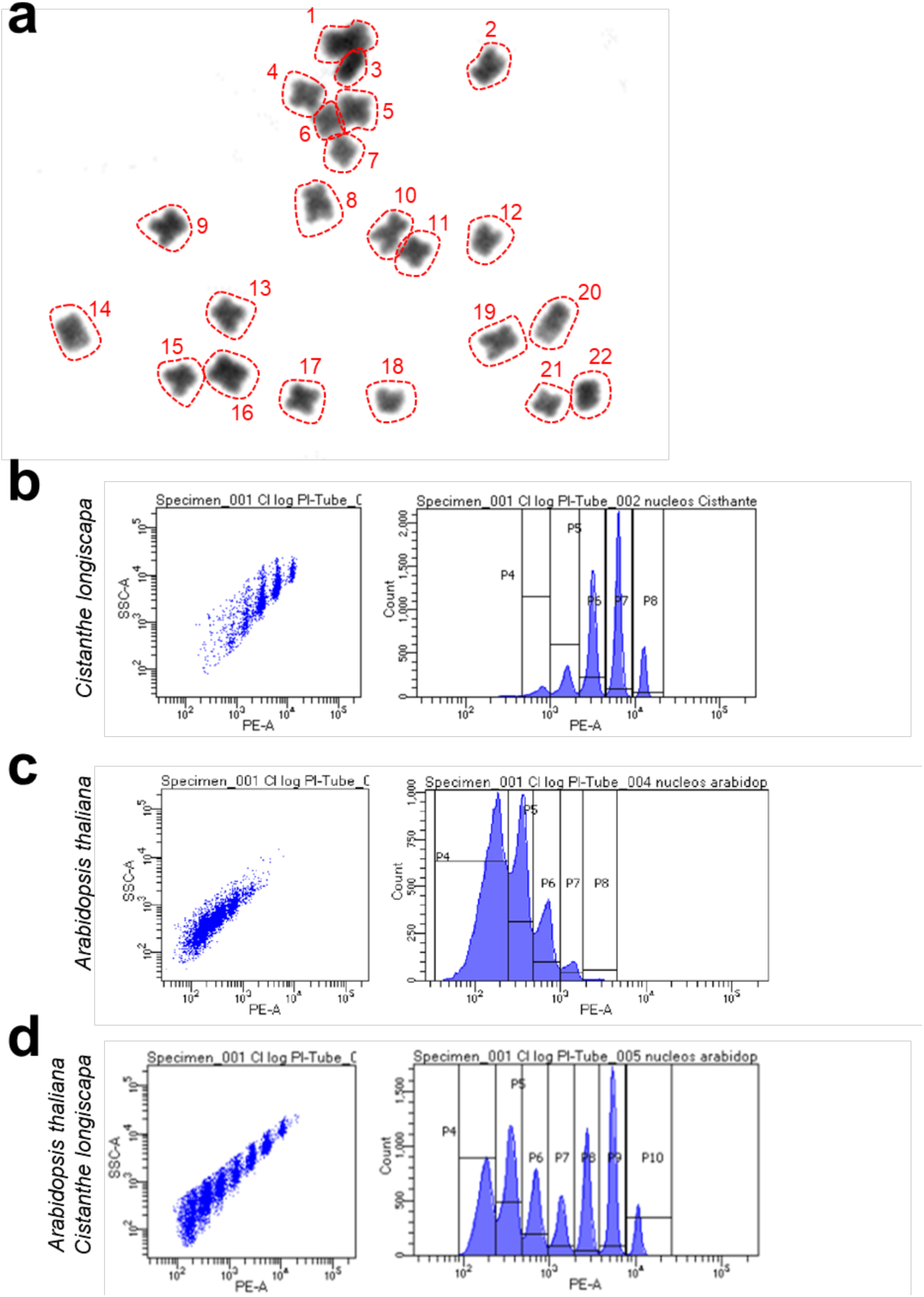
Karyotype analysis and genome size of *Cistanthe longiscapa*. **a)** Karyotyping in root cells of *Cistanthe longiscapa*. A diploid set of 2n= 22 chromosomes was determined in *C. longiscapa*. Individual chromosomes are circled by dotted red lines, and numbered. **b-d)** Genome size estimation by flow cytometry. Isolated nuclei extracted from **b)** *Cistanthe longiscapa*, **c)** *Arabidopsis thaliana* or **d)** a mix of both species were stained with propidium iodide (PI) and subjected to flow cytometry. The side-scattered light (SSC-A; left) and the PI stained histogram (right) are indicated for each sample. The experiment was performed three times (biological replicates) with similar results.

**Fig. S6:**
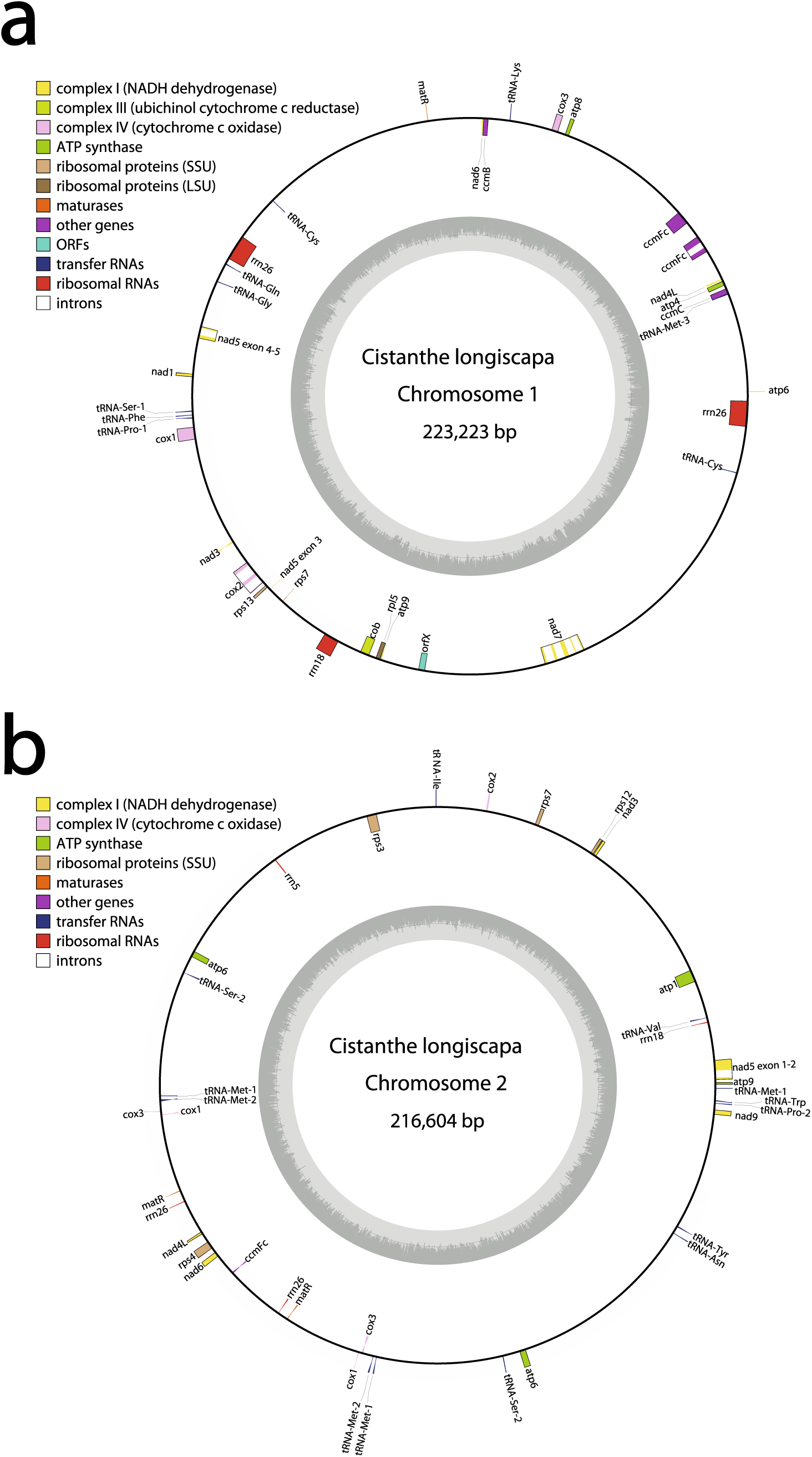
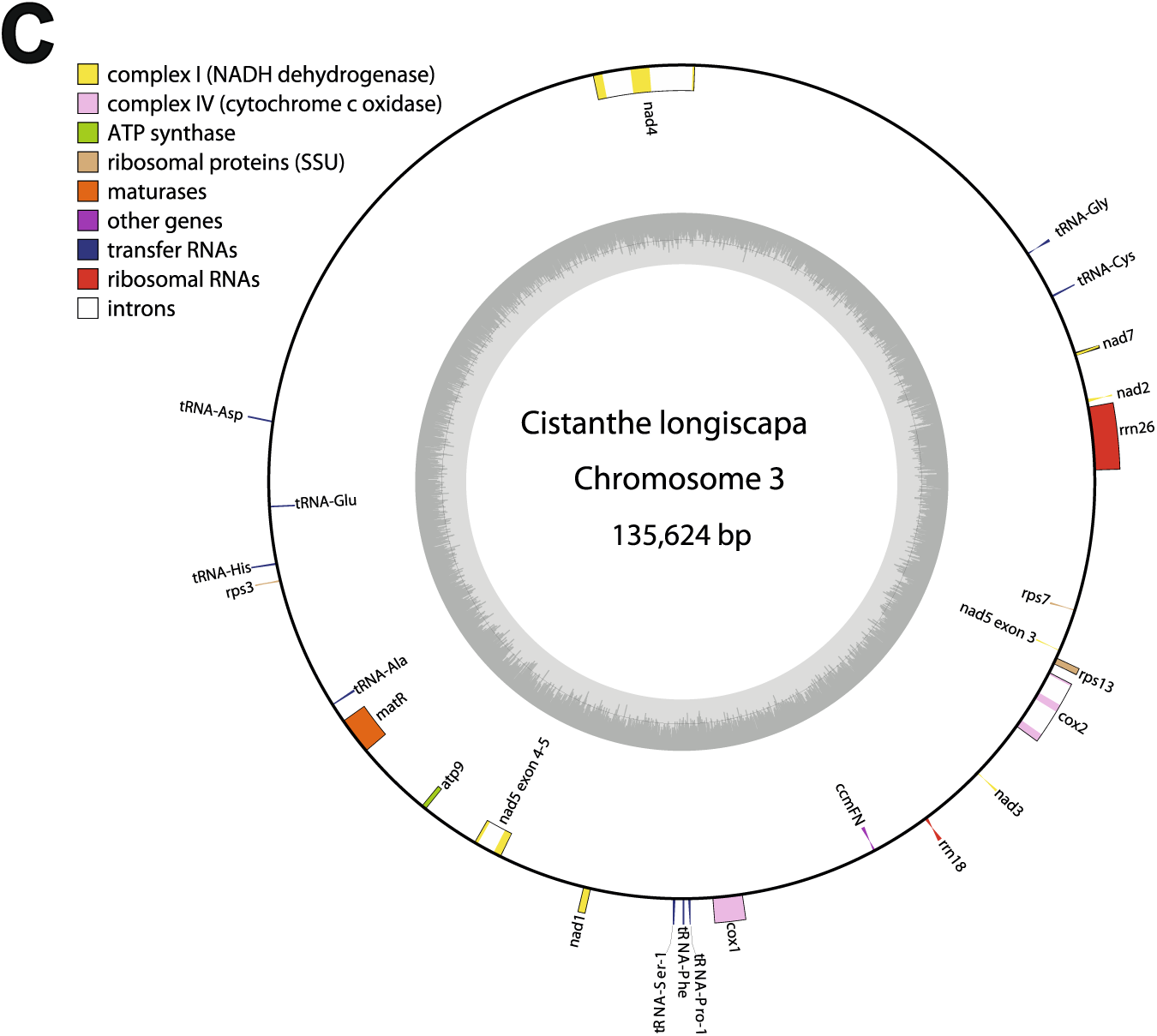
Mitochondrial genome of *Cistanthe longiscapa*. Three different subgenomic circles were identified for the mitochondrial genome of *C. longiscapa*, denoted as **a)** mitochondrial chromosome 1 (223,223 bp), **b)** mitochondrial chromosome 2 (216,604 bp) and **c)** mitochondrial chromosome 3 (135,624 bp).

**Fig. S7:**
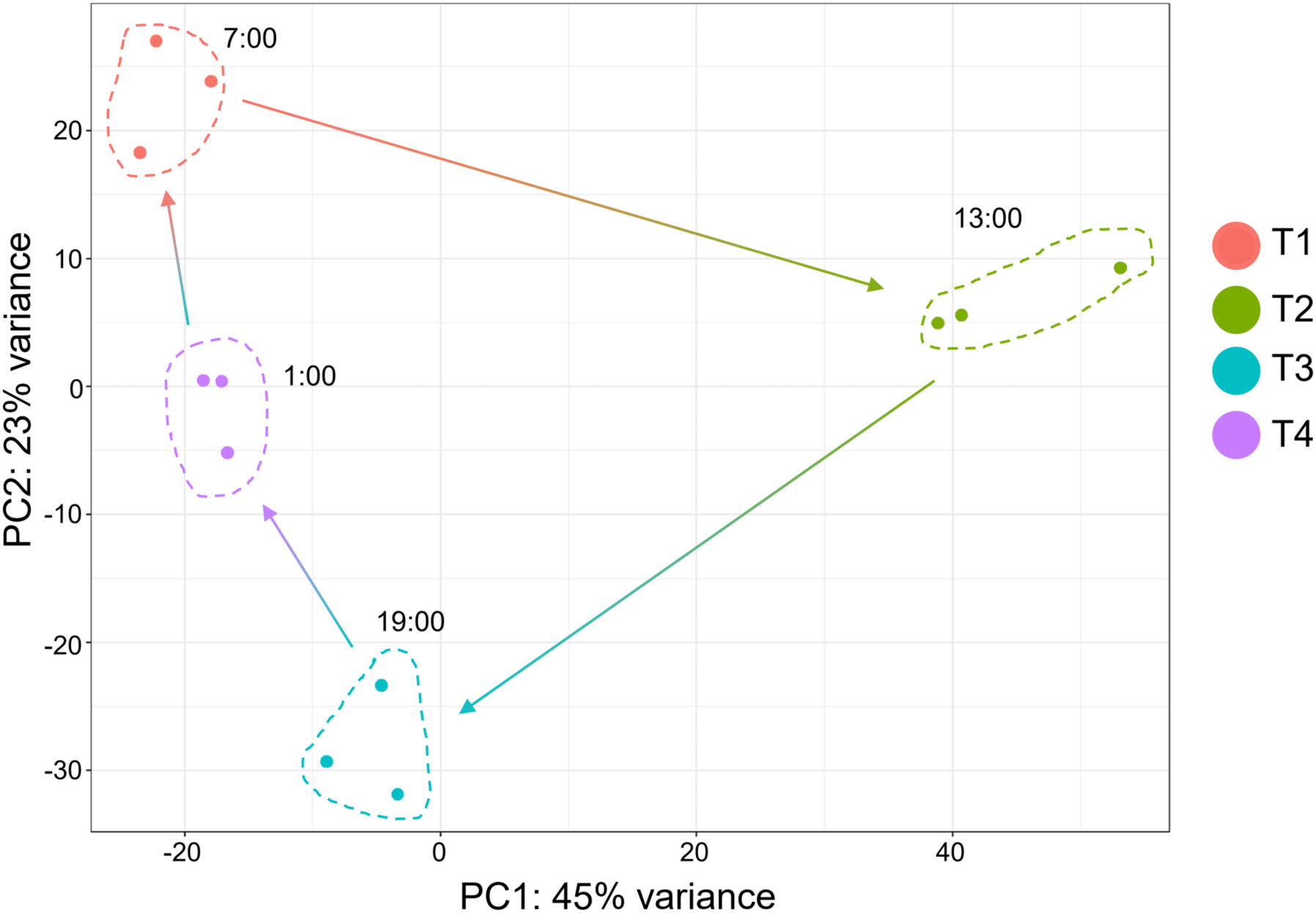
Principal Component Analysis (PCA) showing the distribution and clustering of the individual sample groups. The arrows indicate the temporal succession of the samples in chronological order starting at 7:00 (timepoint T1).

**Fig. S8.**
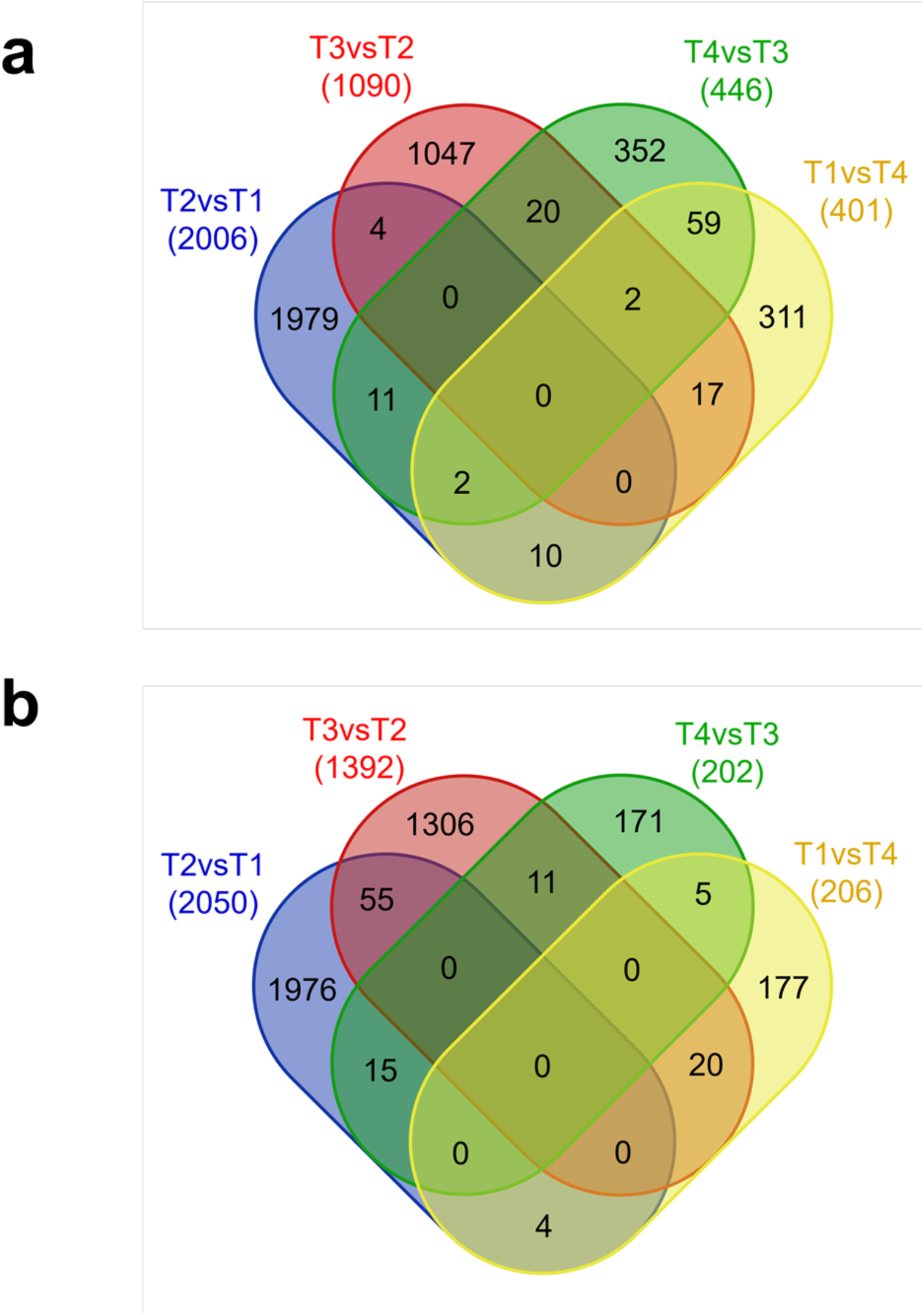
Venn diagram of differentially expressed genes. The diagrams show the distribution of the DEGs showing significant changes in expression in at least one timepoint. The numbers of **a)** overexpressed genes and **b)** downregulated genes are given for each pairwise comparison. The diagrams were constructed with the web-based tool from https://bioinformatics.psb.ugent.be/webtools/Venn/.

**Table S1:** Coordinates of sampling sites.

| Location ID | Coordinates S | Coordinates W | MSL<sup>a</sup> Elevation |
| --- | --- | --- | --- |
| Location 1 | 27°38'01.6" | 70°27'46.6" | 156 |
| Location 2 | 27°56'36.8" | 70°33'25.9" | 453 |
| Location 3 | 28°03'55.1" | 70°35'26.8" | 516 |
| Location 4 | 28°26'22.1" | 70°43'55.6" | 466 |
| Location 5 | 27°33'20.9" | 70°51'10.3" | 170 |

<sup>a</sup> Mean sea level

**Table S2:** List of antibodies employed in this study.

| **Antibody** | **Company / Cat.No.** | **Dilution** |
| --- | --- | --- |
| anti-PsaA | Agrisera / AS06 172 | 1:1000 |
| Anti-PsaB | Agrisera/ AS10 695 | 1:1000 |
| anti-PSAD | Agrisera / AS09 461 | 1:1000 |
| anti-PetA | Agrisera / AS20 4377 | 1:1000 |
| anti-PetB | Agrisera / AS18 4169 | 1:1000 |
| Anti-PetD | Schwenkert et.al., 2007 (Schwenkert et al., 2007) | 1:1000 |
| anti-PsbD | Agrisera / AS06 146 | 1:1000 |
| anti-PsbA | Agrisera / AS10 704 | 1:1000 |
| anti-AtpB | Agrisera / AS05 085 | 1:5000 |
| anti-ATPC | Agrisera / AS08 312 | 1:1000 |
| anti-NdhB | Agrisera / AS16 4064 | 1:1000 |
| anti-LHCA1 | Agrisera / AS01 005 | 1:1000 |
| anti-LHCB2 | Agrisera / AS01 003 | 1:1000 |

**Table S3:** General characteristics of the libraries constructed for sequencing.

| Sequencing runs | Sequencing platform | Libraries | Number of reads | Total base pairs (Mbp) | Mean read length (bp) |
| --- | --- | --- | --- | --- | --- |
| Long reads of *C. longiscapa* genome<sup>a</sup> | PacBio RS II | 25 | 9,738,492 | 89,669.50 | 9,208 |
| Short reads of *C. longiscapa* genome<sup>a</sup> | NovaSeq6000 | 1 | 353,802,412 | 53,424.16 | 151 |
| IsoSeq of *C. longiscapa* | PacBio RS II | 2 | 2,591,616 | 4,964.40 | 1,915 |
| RNAseq of *C. longiscapa* tissues<sup>a</sup> | HiSeq2500 | 11 | 370,950,346 | 37,465.98 | 101 |
| RNAseq of *C. longiscapa* leaf each 6 hours | HiSeq4000 | 12 | 715,693,450 | 72,285.04 | 101 |
| Short reads of *C. longiscapa* individuals for Genetic diversity experiments | HiSeq4000 | 12 | 1,163,305,046 | 117,493.80 | 101 |

<sup>a</sup>Obtained from the same individual

**Table S4:**
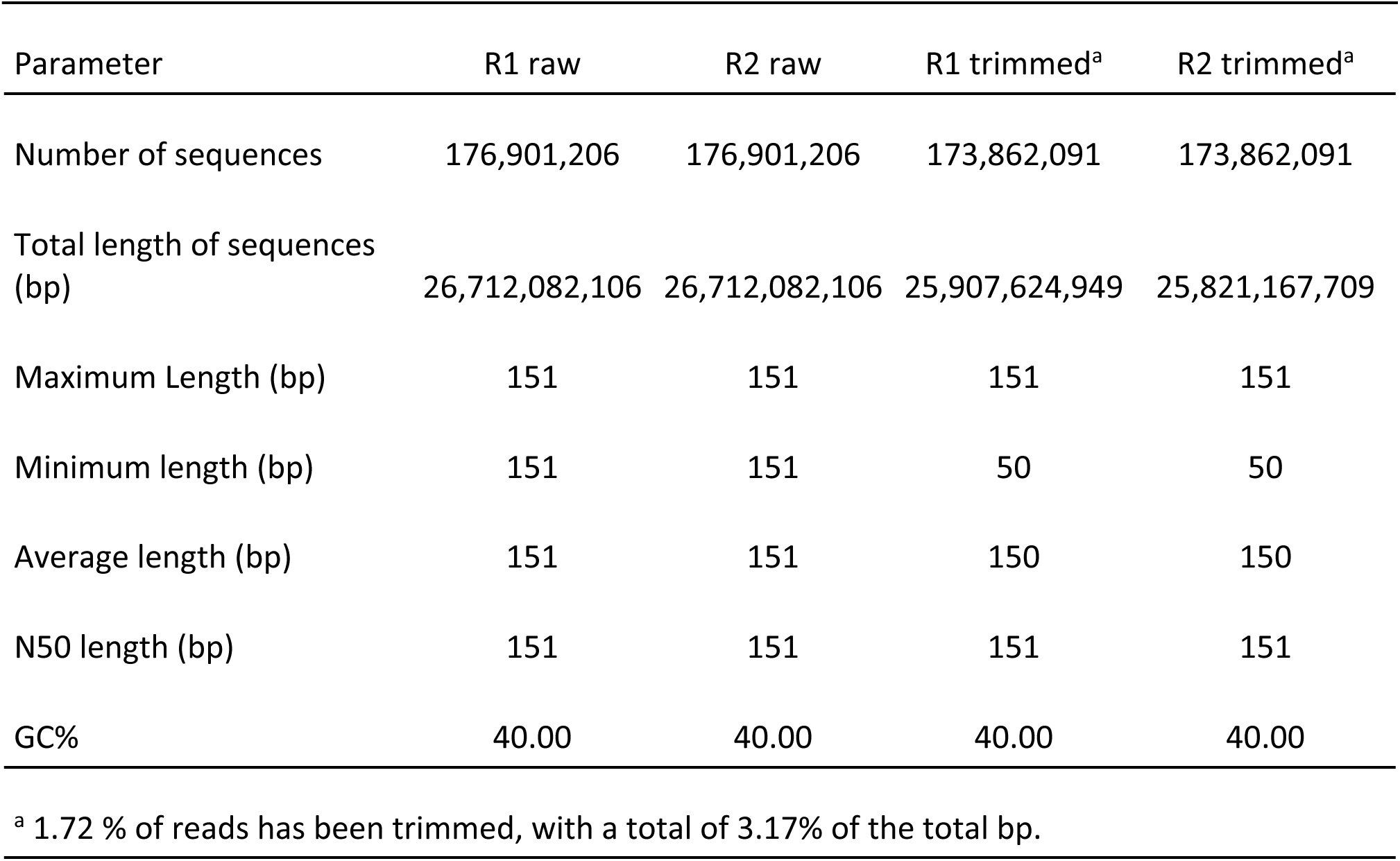
Trimming of short reads of the *Cistanthe longiscapa* genome library for genome assembly.

**Supplementary Table S5:**
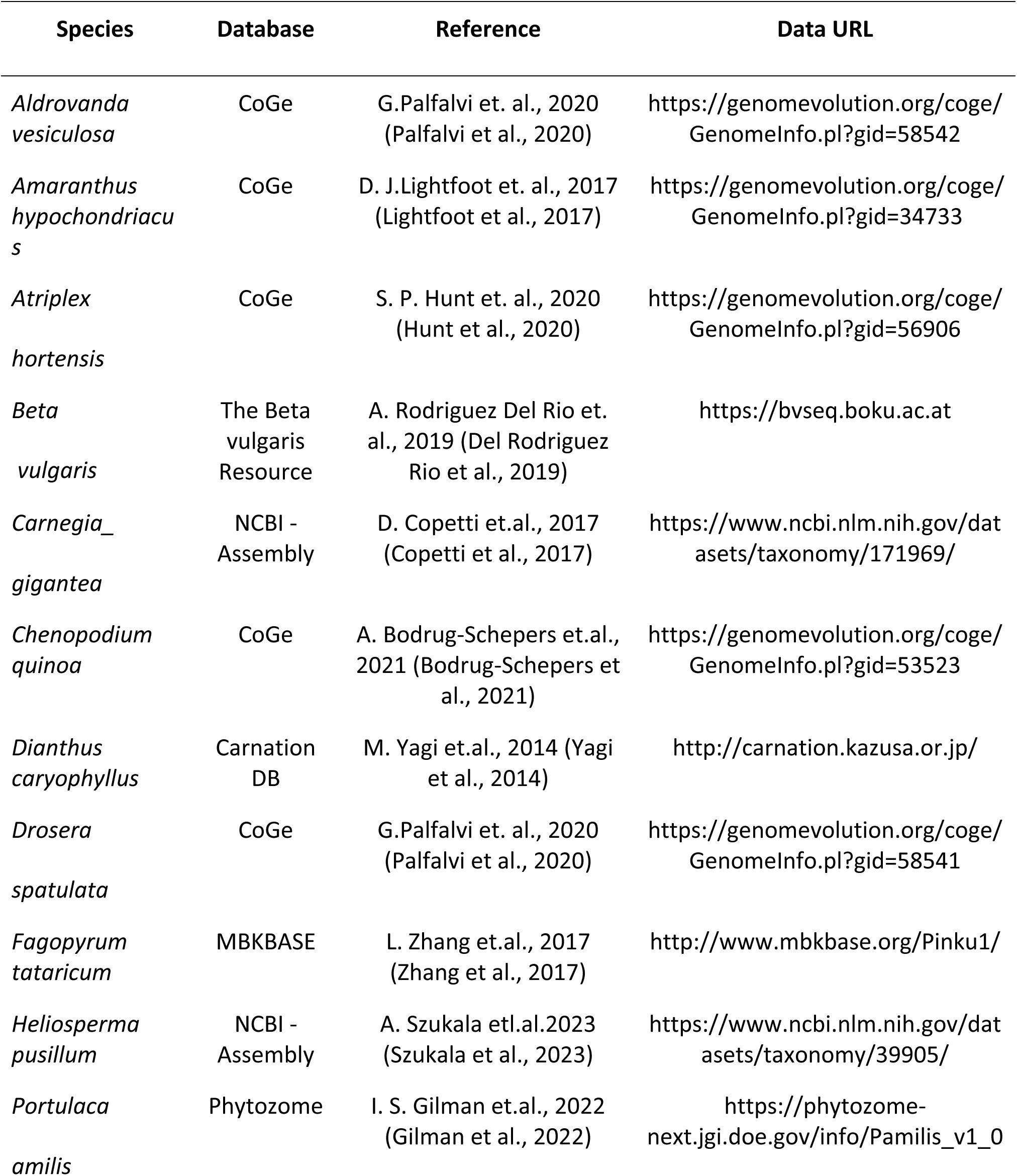
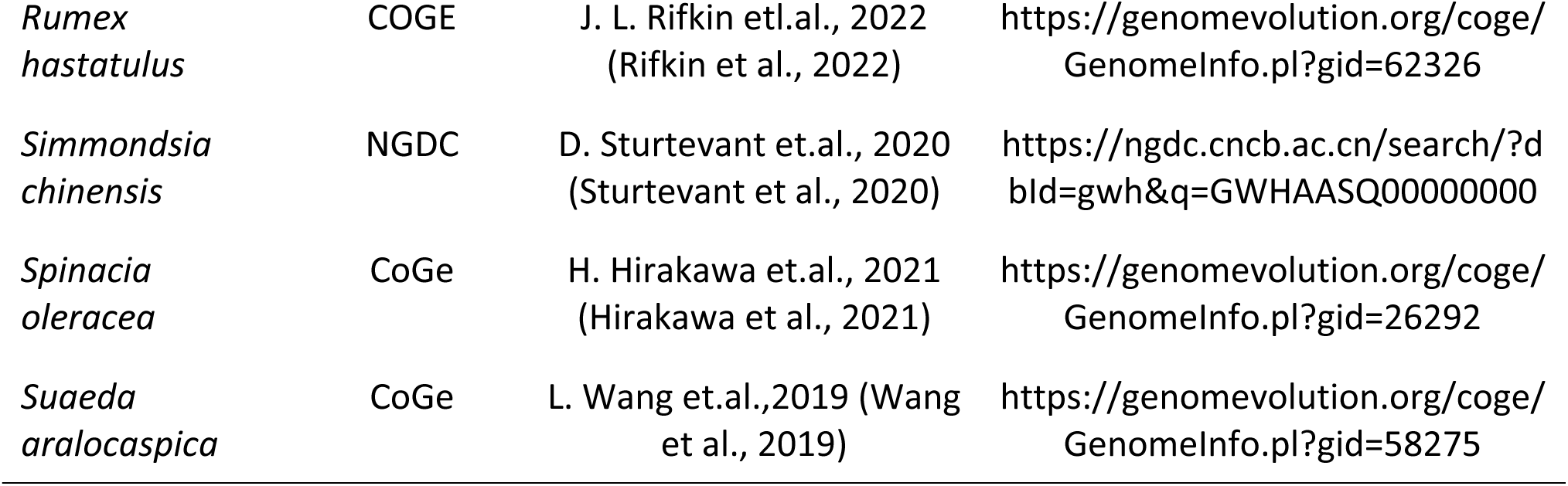
Caryophyllales genomes employed for the nuclear genome assembly.

**Table S6:** *Cistanthe longiscapa* genome assembly and scaffolding statistics.

| **Feature** | **Assembly<sup>a</sup>** | **Scaffolding<sup>b</sup>** |
| --- | --- | --- |
| Number of sequences | 7.586 | 4,831 |
| Total length of sequences (bp) | 786,756,001 | 530,268,464 |
| N25 length (bp) | 263.024 | 10.679.121 |
| N50 length (bp) | 186,054 | 4,743,387 |
| N75 length (bp) | 83.639 | 2.484.445 |
| L50 length (bp) | 27 | 11 |
| GC% | 39.00 | 38.39 |
| N's /100 kbp | 0.10 | 21.64 |
| N total | 0 | 10,076,685 |

<sup>a</sup>Assembly only from Masurca output
<sup>b</sup>Scaffolding output from RagTag

**Table S7:**
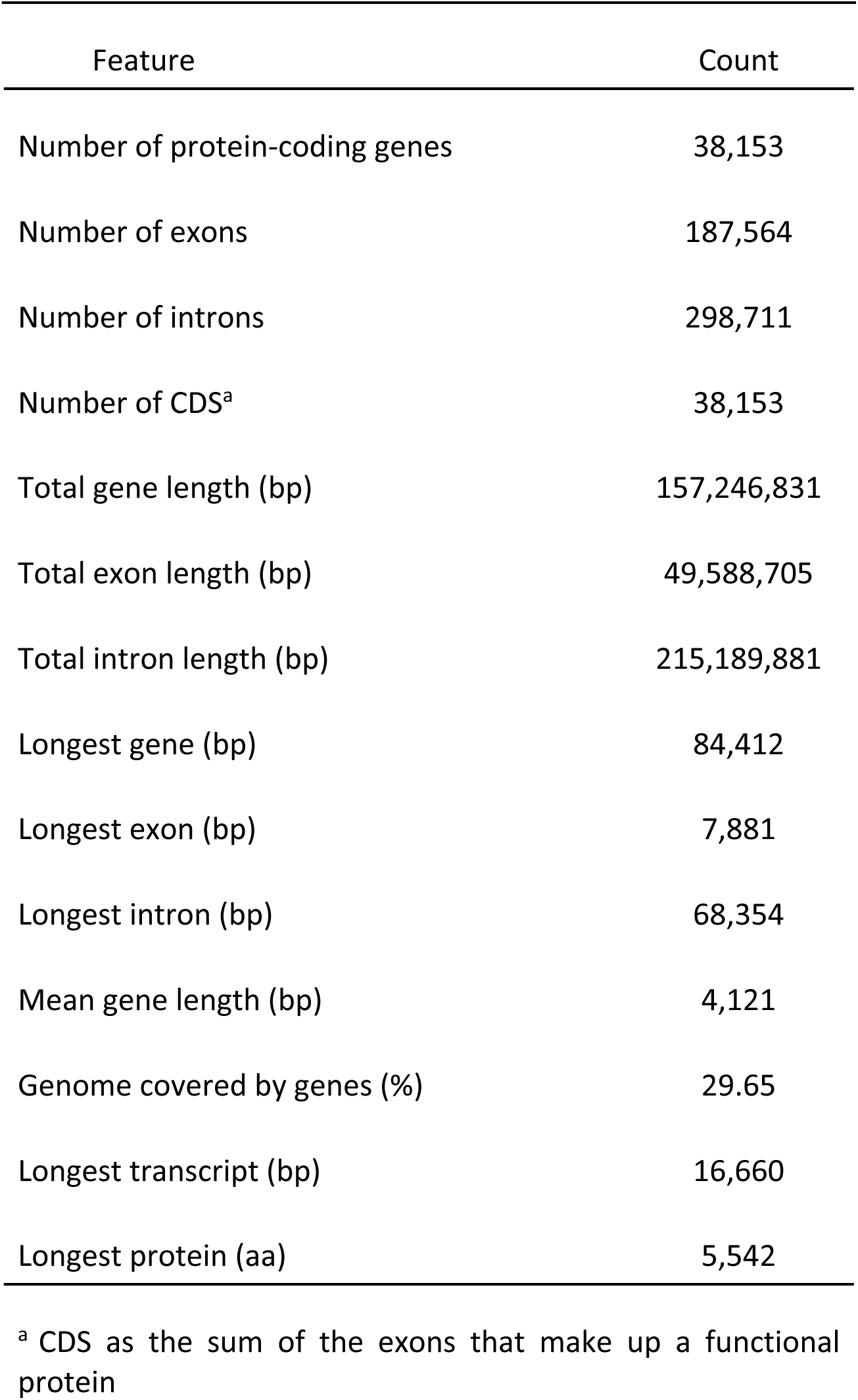
Structural characteristics of the *Cistanthe longiscapa* genome.

| Feature | Count |
| --- | --- |
| Number of protein-coding genes | 38,153 |
| Number of exons | 187,564 |
| Number of introns | 298,711 |
| Number of CDS<sup>a</sup> | 38,153 |
| Total gene length (bp) | 157,246,831 |
| Total exon length (bp) | 49,588,705 |
| Total intron length (bp) | 215,189,881 |
| Longest gene (bp) | 84,412 |
| Longest exon (bp) | 7,881 |
| Longest intron (bp) | 68,354 |
| Mean gene length (bp) | 4,121 |
| Genome covered by genes (%) | 29.65 |
| Longest transcript (bp) | 16,660 |
| Longest protein (aa) | 5,542 |

<sup>a</sup> CDS as the sum of the exons that make up a functional protein

**Table S8:** Benchmarking Universal Single-Copy Orthologs (BUSCO) results for genome scaffolding and genome annotation.

|  | Genome scaffolding | Genome annotation |
| --- | --- | --- |
| BUSCO | Embryophyta (%)<sup>a</sup> | Embryophyta<sup>a</sup> (%) |
| Complete BUSCO | 98.0 | 94.1 |
| Complete and single-copy BUSCO | 87.5 | 84.9 |
| Complete and duplicated BUSCO | 10.5 | 9.2 |
| Fragmented BUSCO | 0.6 | 3.2 |
| Missing BUSCO | 1.4 | 2.7 |

<sup>a</sup> Version 10 of the BUSCO database was used.

**Table S9:** Repetitive elements present in the genome of *Cistanthe longiscapa*.

| Class | Family | Count | Masked (bp)<sup>a</sup> | Masked (%)<sup>a</sup> |
| --- | --- | --- | --- | --- |
| DNA | CMC-EnSpm | 14,461 | 6,872,836 | 1.3 |
|  | Ginger | 6 | 1,990 | 0 |
|  | MULE-MuDR | 6,898 | 4,267,384 | 0.81 |
|  | Maverick | 610 | 136,042 | 0.03 |
|  | Novosib | 41 | 4,029 | 0 |
|  | PIF-Harbinger | 1,779 | 565,830 | 0.11 |
|  | Sola-1 | 1 | 134 | 0 |
|  | TcMar-Stowaway | 3,048 | 650,418 | 0.12 |
|  | hAT | 1,324 | 1,066,961 | 0.2 |
|  | hAT-Ac | 13,359 | 4,182,407 | 0.79 |
|  | hAT-Charlie | 221 | 77,791 | 0.01 |
|  | hAT-Tag1 | 5,278 | 1,995,452 | 0.38 |
|  | hAT-Tip100 | 1,904 | 645,909 | 0.12 |
| LINE | L1 | 19,550 | 17,187,348 | 3.24 |
|  | L1-Tx1 | 127 | 28,896 | 0.01 |
|  | L2 | 782 | 273,892 | 0.05 |
|  | Penelope | 3 | 408 | 0 |
|  | R2-NeSL | 1,117 | 1,182,643 | 0.22 |
|  | RTE-BovB | 7,622 | 1,537,263 | 0.29 |
| LTR | Cassandra | 1,737 | 648,014 | 0.12 |
|  | Caulimovirus | 132 | 110,566 | 0.02 |
|  |  | 23,357 | 17,230,218 | 3.25 |
|  | Copia |  |  |  |
|  | ERV1 | 1,690 | 434,602 | 0.08 |
|  | ERVK | 52 | 3,010 | 0 |
|  | Gypsy | 43,404 | 39,448,861 | 7.44 |
|  | Pao | 354 | 110,347 | 0.02 |
|  | unkown | 57,790 | 51,400,446 | 9.7 |
| RC | Helitron | 4,458 | 1,607,113 | 0.3 |
|  | Helitron-2 | 5 | 337 | 0 |
| SINE | RTE | 3,187 | 301,615 | 0.06 |
|  | tRNA-Deu-L2 | 35 | 2,600 | 0 |
|  | tRNA-RTE | 14,775 | 1,516,300 | 0.29 |
| Unknown | -- | 233,318 | 75,384,806 | 14.22 |
| Total interspersed | -- | 476,119 | 231,445,745 | 43.66 |
| Low_complexity | -- | 243 | 39,065 | 0.01 |
| Satellite | -- | 867 | 356,699 | 0.07 |
| Simple_repeat | -- | 16,413 | 4,215,582 | 0.8 |
| Total | -- | 493,642 | 236,057,091 | 44.52 |

<sup>a</sup>Fraction of the reference genome sequence replaced with Ns for masking of repetitive elements.

**Table S10:** Number of annotated functional protein-coding genes.

| **Protein coding genes annotation** | **annotated**<br>**(count)** | **annotated**<br>**(%)** |
| --- | --- | --- |
| Swissprot viridiplantae homology | 20,837 | 54.61 |
| Araport11 homology | 21,259 | 55.72 |
| UniProt viridiplantae | 20,837 | 54.61 |
| Non-redundant Blast | 15,913 | 41.71 |
| With EggNOG mapper 5.0 | 16,779 | 43.98 |
| Mercator Biological process | 21,076 | 55.24 |
| Mercator Description | 19,726 | 51.70 |
| MAKER | 22,195 | 58.17 |
| Only GO term from EggNOG 5.0 | 13,703 | 35.92 |
| Only KO term from EggNOG mapper 5.0 | 13,517 | 35.43 |
| With Mercator4/mapman | 21,076 | 55.24 |
| Total annotated<sup>b</sup> | 32,714 | 85.74 |
| Non-annotated | 5,439 | 14.26 |

<sup>a</sup> Annotated with all InterProScan databases<sup>b</sup> Annotated with at least one annotation database

**Table S11:** Overview of the total number of SNP and indel polymorphisms identified across the genome of *Cistanthe longiscapa* specimens from various geographic locations.

| Sample | SNPs |  |  |  | InDel |  |  |
| --- | --- | --- | --- | --- | --- | --- | --- |
|  | Total | Het | Hom | TS/TV | Total | Het | Hom |
| Reference genome | 3,217,020 | 3,213,377 | 3,643 | 1.46 | 629,499 | 626,208 | 3,291 |
| Loc2_WGS_BR1 | 3,140,888 | 1,359,532 | 1,781,356 | 1.44 | 538,182 | 251,146 | 287,036 |
| Loc2_WGS_BR2 | 3,299,055 | 1,463,697 | 1,835,358 | 1.44 | 571,015 | 273,437 | 297,578 |
| Loc2_WGS_BR3 | 3,288,030 | 1,475,222 | 1,812,808 | 1.43 | 571,888 | 278,453 | 293,435 |
| Loc3_WGS_BR1 | 2,940,193 | 1,468,348 | 1,471,845 | 1.43 | 495,496 | 264,420 | 231,076 |
| Loc3_WGS_BR2 | 3,306,670 | 1,444,142 | 1,862,528 | 1.44 | 570,808 | 268,792 | 302,016 |
| Loc3_WGS_BR3 | 3,261,768 | 1,422,295 | 1,839,473 | 1.43 | 571,550 | 271,189 | 300,361 |
| Loc5_WGS_G1_BR1 | 3,362,005 | 1,299,764 | 2,062,241 | 1.43 | 598,315 | 250,614 | 347,701 |
| Loc5_WGS_G1_BR2 | 3,324,391 | 1,262,388 | 2,062,003 | 1.43 | 591,721 | 243,992 | 347,729 |
| Loc5_WGS_G1_BR3 | 3,224,568 | 1,197,477 | 2,027,091 | 1.43 | 574,263 | 231,033 | 343,230 |
| Loc5_WGS_G2_BR1 | 2,143,721 | 564,723 | 1,578,998 | 1.43 | 367,908 | 105,787 | 262,121 |
| Loc5_WGS_G2_BR2 | 3,129,729 | 1,122,766 | 2,006,963 | 1.43 | 556,658 | 216,599 | 340,059 |
| Loc5_WGS_G2_BR3 | 2,593,483 | 829,455 | 1,764,028 | 1.43 | 449,148 | 153,307 | 295,841 |

**Table S12:** Analysis of polymorphisms identified in coastal and inland populations of *Cistanthe longiscapa*.

| Description | Sample | SNPs |  |  |  |
| --- | --- | --- | --- | --- | --- |
|  |  | Total | Het | Hom | TS/TV |
| **Conserved across locations** | Loc5_WGS_G1 | 253,170 | 37,123 | 216,047 | 1.43 |
|  | Loc5_WGS_G2 | 145,943 | 13,143 | 132,800 | 1.41 |
|  | Loc2_WGS | 122,881 | 20,632 | 102,249 | 1.43 |
|  | Loc3_WGS | 165,117 | 36,081 | 129,036 | 1.43 |
| **Conserved in coastal** | Intersection of SNPs between Loc5_WGS_G1 and Loc5_WGS_G2 | 48,191 | 1,402 | 46,789 | 1.38 |
| **Conserved in inland** | Intersection of SNPs between Loc2_WGS and Loc3_WGS | 26,329 | 859 | 25,470 | 1.40 |
| **Common in all samples** | Intersection of SNPs between coastal + inland | 4,092 | 51 | 4,041 | 1.29 |
| **Unique in coastal** | Unique coastal | 44,099 | 1,351 | 42,748 | 1.39 |
| **Unique in inland** | Unique inland | 22,237 | 808 | 21,429 | 1.42 |

**Table S13.**
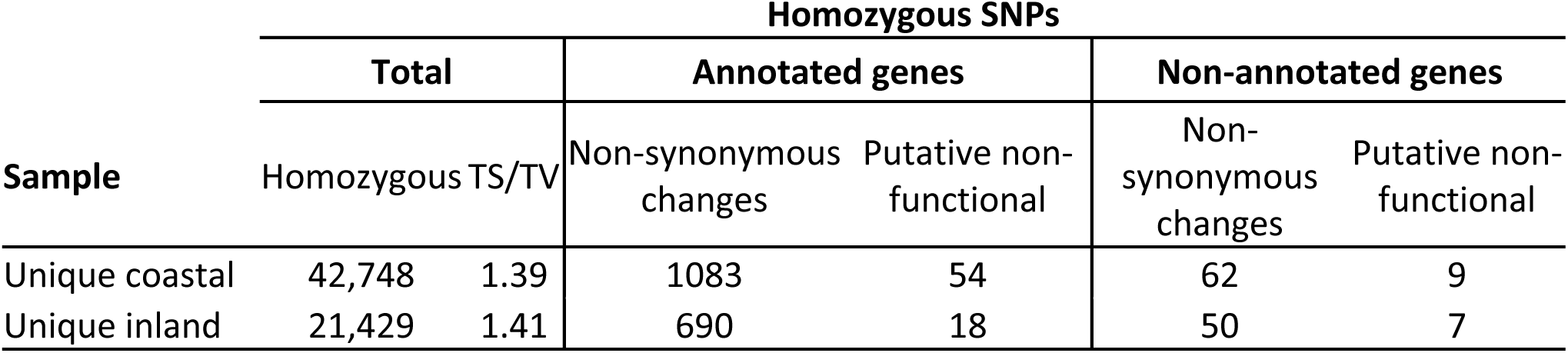
Overview of homozygous polymorphisms affecting gene functions in coastal and inland populations of *Cistanthe longiscapa*.

## Supplemental Methods

**Methods S1: Settings for Imaging Pulse-Amplitude Modulation (Imaging-PAM) analysis.** We used the following settings for data collection: Gain=1, Set Damping=2, Measur.light=3, means Freq=1. The induction curves and the recovery for *C. longiscapa* and *A. hypochondriacus* were conducted using the following program: 33 μmol photons m^−2^ s^−1^ 150 s; 55 μmol photons m^−2^ s^−1^ 150 s, 80 μmol photons m^−2^ s^−1^ 120 s, 108 μmol photons m^−2^ s^−1^ 100 s, 144 μmol photons m^−2^ s^−1^ 100 s, 185 μmol photons m^−2^ s^−1^ 200 s, 230 μmol photons m^−2^ s^−1^ 100 s, 280 μmol photons m^−2^ s^−1^ 100 s, 335 μmol photons m^−2^ s^−1^ 100 s, 530 μmol photons m^−2^ s^−1^ 100 s, 700 μmol photons m^−2^ s^−1^ 100 s, 920 μmol photons m^−2^ s^−1^ 100 s, 1,160 μmol photons m^−2^ s^−1^ 100 s, 0 μmol photons m^−2^ s^−1^ 10 s, 0 μmol photons m^−2^ s^−1^ 10 s, 0 μmol photons m^−2^ s^−1^ 10 s, 0 μmol photons m^−2^ s^−1^ 50 s, 0 μmol photons m^−2^ s^−1^ 50 s, 0 μmol photons m^−2^ s^−1^ 100 s and 0 μmol photons m^−2^ s^−1^ 100 s. The induction curves and the recovery for *C. longiscapa* and *C. grandiflora* were analyzed using the following program: 33 μmol photons m^−2^ s^−1^ 150 s; 55 μmol photons m^−2^ s^−1^ 150 s, 80 μmol photons m^−2^ s^−1^ 120 s, 108 μmol photons m^−2^ s^−1^ 100 s, 144 μmol photons m^−2^ s^−1^ 100 s, 185 μmol photons m^−2^ s^−1^ 200 s, 230 μmol photons m^−2^ s^−1^ 100 s, 280 μmol photons m^−2^ s^−1^ 100 s, 530 μmol photons m^−2^ s^−1^ 100 s, 700 μmol photons m^−2^ s^−1^ 100 s, 920 μmol photons m^−2^ s^−1^ 100 s, 1,160 μmol photons m^−2^ s^−1^ 100 s, 0 μmol photons m^−2^ s^−1^ 10 s, 0 μmol photons m^−2^ s^−1^ 10 s, 0 μmol photons m^−2^ s^−1^ 10 s, 0 μmol photons m^−2^ s^−1^ 50 s, 0 μmol photons m^−2^ s^−1^ 50 s, 0 μmol photons m^−2^ s^−1^ 100 s and 0 μmol photons m^−2^ s^−1^ 100 s.

**Methods S2: Genome assembly of C. longiscapa.** To assemble a pre-processed library based on Illumina reads, the Genome_NS_T01 library was trimmed by removing low quality 3’ borders, sequencing adaptors and reads with less than 35 bp using the Trim_galore v.0.6.5 software with default settings (https://github.com/FelixKrueger/TrimGalore). Plastid reads were removed from the trimmed library by alignment to the *C. longiscapa* reference chloroplast genome (Stoll et al., 2017) using the Bowtie2 v.2.3.5.1 software (Langmead and Salzberg, 2012) employing the default settings except for -p defined as 40 and the -very-sensitive-local option. Finally, the reads aligned to the plastid genome were sorted by coordinate, and all aligned reads were removed using the Samtools v.1.10 software package (Li et al., 2009). The unaligned reads were saved as a fastq file for hybrid correction of the PacBio libraries and assembly of a hybrid reference genome. For the hybrid correction of the PacBio library, the 25 libraries from the PacBio genome sequencing (Genome_PB_T01 to Genome_PB_T25) were concatenated into a single fasta file. The resulting file was corrected using the LORDEC v.0.9 software (Salmela and Rivals, 2014) with the previously constructed pre-processed Illumina library (Genome_NS_T01) as an additional input and using default settings except for “[-c] -s 3” with a k-mer value of 21, resulting in a fasta file with corrected PacBio data.

For the assembly of a hybrid reference genome, the raw PacBio libraries obtained from *C. longiscapa* (libraries Genome_PB_T01 to Genome_PB_T25) were concatenated into a single fasta file. The resulting file was assembled and combined with the plastid-free Illumina pre-processed reads using the software MaSuRCA v.4.0.8 (Zimin et al., 2017) with the option “FLYE_ASSEMBLY=1” in the configuration file. The resulting draft hybrid genome was analyzed using the software Purge_Dups v.1.2.5 (Guan et al., 2020) to confirm unique contigs, resulting in a refined version of the draft genome assembly, and hybrid scaffold with the concatenated PacBio data using LongStitch v.1.0.2 (Coombe et al., 2021) with the option “tigmint-ntLink-arks”. The resulting scaffold reference genome was refined in two steps. First, the gaps were removed using the software TGS-GapCloser v.1.0.1(Xu et al., 2020) with the Lordec corrected PacBio data as input with the option “--ne”. The erroneous consensus base pairs were then corrected using the POLCA software (Zimin and Salzberg, 2020) included as part of the MaSuRCA pipeline with default settings, resulting in a refined version of the reference genome scaffolds. The refined genome scaffold was rearranged using the software RagTag v.2.1.0 using the genome assembly of *Portulaca amilis* (Gilman et al., 2022), *Selenicereus undatus* (Zheng et al., 2021) and *Talinum fruticosum* (NCBI Accession: PRJNA659383; ID: 659383) as references, obtaining the final version of the reference genome. The genome assembly and scaffolding were validated using BUSCO v.5.2.2 software (Manni et al., 2021) to assess the completeness of the PacBio raw data, the long-read primary assembly, and the hybrid scaffolding. Additionally, each library was aligned to the genome scaffold using BWA mem v.0.0.17 (r1188) for Illumina libraries (arXiv:1303.3997v2), and Minimap2 v.2.24(r1122) for PacBio concatenated libraries (Li, 2018).

**Methods S3: Genome annotation of *C. longiscapa.*** To annotate the *C. longiscapa* genome we used the MAKER-P v.3.01.03 software (Campbell et al., 2014b). To annotate repetitive elements and create a custom library for RepeatMasker v.4.1.2, we followed the recommendations of the authors (Campbell et al., 2014a) and the software wiki (Campbell et al., 2014b). The resulting file was converted to the annotated GFF3 format using RepeatCraft v1.0 (Wong and Simakov, 2019). For ab-initio prediction, we followed the recommendation of the authors and the software wiki (Campbell et al., 2014a), and used the fasta version of SwissProt-uniprot viridiplantae retrieved in December 2021 as a protein database in addition to the complete proteome of 15 *Caryophyllales* genomes.

As part of the MAKER-P annotation, a total of 25 *C. longiscapa* transcriptome libraries were used as input data. These included 11 Illumina RNA-Seq datasets from *C. longiscapa* tissue samples, 12 Illumina RNA-Seq datasets from leaf samples collected at six-hour intervals, and 2 Iso-Seq libraries. Reads were concatenated into a single FASTA archive and filtered to remove repeat sequences using USEARCH v.11.0.667 (Edgar, 2010) with the -fastx_uniques option. The annotation included training data from SNAP v.2006-07-28 (Korf, 2004), Augustus v.3.2.3 (Stanke et al., 2008) and GeneMark-ES v.4.69_lic (Ter-Hovhannisyan et al., 2008) predictions as well as the predicted proteins and transcript data mentioned above. The results of the annotation are summarized in Supplementary Table S5, considering only predictions with an AED < 0.75 after validation by EvidenceModeler v.1.1.0 (Haas et al., 2008). Completeness of annotation was assessed using BUSCO v.5.2.2 software (Manni et al., 2021) (Supplementary Table S6). Finally, functional annotation was performed by BLASTp alignment of predicted proteins-coding genes to the SwissProt-viriplantae database and InterProScan to the Pfam database. To complement the annotation, we also used the online platforms Mercator4/Mapman (https://plabipd.de/portal/mercator4, November 2022), eGGNog mapper v.2 with eGGNog v.5.0 Database (http://eggnog5.embl.de/#/app/home, July 2022) and KASS (https://www.genome.jp/kegg/kaas/, November 2022).

**Methods S4: Comparative genomics.** To build the ultrametric tree, the proteomes of 15 Caryophyllales species were obtained from different repositories. The sequences from the repositories, as well as those from *A. thaliana* and *C. longiscapa* were used as input in Orthofinder v.2.5.5 software (Emms and Kelly, 2019) with the option “-M msa”. The resulting files consisted of the rooted species tree with node names, and the number of orthologs for each orthogroup for each species proteome. The rooted species tree with node names file was used to generate the ultrametric tree with the PATHd8 v1.0 software (Britton et al., 2007) using three calibration points obtained from www.timetree.org (Kumar et al., 2022) for *A. thaliana* – *Fagopyrum tataricum* (125 MYA), *Portulaca amilis* – *Chenopodium quinoa* (90 MYA), and *Fagopyrum tataricum* – *Beta vulgaris* (106 MYA). The expansion/contraction of orthologous groups was computed with the software Cafe v.5.0.0 (Mendes et al., 2021) with a lambda of 0.0051026692207754, using as input the file with the number of orthologs for each orthogroup obtained by Orthofinder and the ultrametric tree obtained from PATHd8.

For the Gene Ontology enrichment analysis, the annotated predicted proteins of the *C. longiscapa* genome were aligned with the complete *A. thaliana* proteome retrieved from SwissProt using BlastP to generate a list of homologous genes. The list was used as input to the “enrichGO” function from the R language package clusterProfiler v3.14.3 (Yu et al., 2012), and filtered to the fourth level of gene ontology terms using the “dropGO” function from the same package. Plots for the enriched terms were generated using the dotplot function of the clusterProfiler package (Yu et al., 2012).

**Methods S5: Short variant calling.** For short variant discovery, the raw data of 12 WGS libraries from three different locations (location 2, 3 and 5; Table S1) were normalized to the library with the lower number of reads (Loc5_Individual4_BR1; 53,074,278 reads after trimming) using the software seqtk v.1.3-r106 with the function “sample”, with a seed of 100 (-s 100) and 53074278 as “ <FRAC>| <NUMBER>” option. The normalized libraries were converted to uBAM format, and short variant discovery (SVD) was performed with GATK v.4.0.9.0 (van der Auwera et al., 2013) using the final scaffolded version of the reference genome. The final 12 VCF files (one for each library) were filtered for QUAL > 100. To obtain the unique or shared SNPs for each VCF file, we used the “isec” package from bcftools v.1.9 (Danecek et al., 2021).The identification of SNPs that generate non/functional proteins was performed using SNPeff (Cingolani et al., 2012).

**Methods S6: Draft assembly of the mitochondrial genome.** Raw data from five libraries of short Illumina reads from location 5 (Loc5_individual2_BR1, Loc5_individual2_BR2, Loc5_individual2_BR3, Loc5_individual4_BR1, Loc5_individual4_BR2; Dataset S1) were used to assemble the mitochondrial genome. The initial assembly was generated using GetOrganelle v1.7.7.0 (Jin et al., 2020). The maximum number of extension rounds were set to 20, and the SPAdes k-mer parameters were set to 21, 45, 65, 85, and 105. The source of the contigs was identified by BLASTN using the Caryophyllales genomes, and filtered by Bandage v0.8.1 (Wick et al., 2015). The annotation of protein-coding genes and rRNAs was performed by the online version of GeSeq v2.03 (Tillich et al., 2017) using the Caryophyllales mitochondrial genes as reference, and later were manually verified. For tRNA gene annotation, we used the GeSeq built-in tRNAscan-SE v2.0.7 (Chan et al., 2021). Maps of the assembled mitochondrial DNA circles were drawn using the OGDRAW tool v1.1.1 (Lohse et al., 2007; Greiner et al., 2019).

**Methods S7: Differential expression analysis and GO annotation.** For quality assessment, the reads obtained from the RNA sequencing were first evaluated using FastQC software v0.11.7 (https://www.bioinformatics.babraham.ac.uk/projects/fastqc/) (Liu et al., 2013). The reads with a base quality of Q≤25 phred 53 and length < 50 bp were filtered, and the adapters were removed using Trim Galore v0.6.5. Finally, the post-filtered reads were analyzed using FastQC as a final quality control prior to alignment. Differential expression analysis was performed by comparing the libraries Location 4_7:00_BR1, Location 4_7:00_BR2, Location 4_7:00_BR3, Location 4_13:00_BR1, Location 4_13:00_BR1, Location 4_13:00_BR1, Location 4_19:00_BR1, Location 4_19:00_BR2, Location 4_19:00_BR3, Location 4_01:00_BR1, Location 4_01:00_BR2 and Location 4_01:00_BR3, all obtained from location 4 (Dataset S1). The filtered reads were mapped to the reference genome using STAR v2.7.10a (Dobin et al., 2013), and were counted using the Rsubread v2.2.2 package (Liao et al., 2019). Differential expression analysis was performed using DESeq2 v1.26.0 (Anders and Huber, 2010). Reads with counts lower than 5 were discarded, and the data were normalized to the libraries obtained at 7:00 using the median of ratios method. Sequences annotated as plastid- and mitochondrial-encoded transcripts were removed, and finally, the data were filtered by fold change (-1>FC>1), using an FDR <0.05.

For GO enrichment annotation of consecutive time points, differentially expressed transcripts up- and down-regulated transcripts obtained in each pairwise comparison (T2/T1, T3/T2, T4/T3 and T1/T4) were analyzed with the Clusterprofiler v3.16.0 R package (Yu et al., 2012), using the Araport11 *Arabidopsis thaliana* annotation as input, and an adjusted *P*-value <0.05.

